# Real-time genomic characterization of pediatric acute leukemia using adaptive sampling

**DOI:** 10.1101/2024.10.11.617690

**Authors:** Julie Geyer, Kofi B. Opoku, John Lin, Lori Ramkissoon, Charles Mullighan, Nickhill Bhakta, Thomas B. Alexander, Jeremy R. Wang

## Abstract

Effective treatment of pediatric acute leukemia is dependent on accurate genomic classification, typically derived from a combination of multiple time-consuming and costly techniques such as flow cytometry, fluorescence *in situ* hybridization (FISH), karyotype analysis, targeted PCR, and microarrays (Arber et al., 2016; Iacobucci & Mullighan, 2017; Narayanan & Weinberg, 2020). We investigated the feasibility of a comprehensive single-assay classification approach using long-read sequencing, with real-time genome target enrichment, to classify chromosomal abnormalities and structural variants characteristic of acute leukemia. We performed whole genome sequencing on DNA from diagnostic peripheral blood or bone marrow for 57 pediatric acute leukemia cases with diverse genomic subtypes. We demonstrated the characterization of known, clinically relevant karyotype abnormalities and structural variants concordant with standard-of-care clinical testing. Subtype-defining genomic alterations were identified in all cases following a maximum of forty-eight hours of sequencing. In 18 cases, we performed real-time analysis – concurrent with sequencing – and identified the driving alteration in as little as fifteen minutes (for karyotype) or up to six hours (for complex structural variants). Whole genome nanopore sequencing with adaptive sampling has the potential to provide genomic classification of acute leukemia specimens with reduced cost and turnaround time compared to the current standard of care.

## Introduction

B-cell acute lymphoblastic leukemia (B-ALL) is the most common type of pediatric cancer. B-ALL shares genomic underpinnings across the age spectrum into late adulthood, albeit with increased relative frequency of high-risk genomic subtypes. Acute myeloid leukemia (AML) accounts for 15-20% of pediatric acute leukemia cases and is defined by large-scale structural variation and small sequence variations, which, similarly to B-ALL, are detectable with whole-genome sequencing (de Rooij, Zwann & van den Heuvel-Eibrink, 2015). Currently, B-ALL and AML cases are classified into clinical genomic subtypes through a cascade of internal and external testing that is complex, costly, and often lacks comprehensiveness. These tests include karyotype analysis, fluorescence *in situ* hybridization (FISH), polymerase chain reaction (PCR) testing, targeted sequencing panels, and/or comprehensive genomic sequencing (Arber, 2016; Iacobucci, 2017; Narayanan, 2020). The "standard-of-care" diagnostic pipeline varies greatly by testing facility, with diagnosis in resource-limited areas being particularly challenging due to limited availability of diagnostic tools (Gupta et al., 2015). The multiple distinct, costly, and laborious techniques, which are necessary to accurately classify pediatric acute leukemia into clinically relevant genomic subtypes, have implications for prognosis, selection of treatment intensity, and precision medicine approaches.

Single assay sequencing-based classification approaches are a potential solution to reduce the complexity of the standard-of-care cascade of tests, minimize costs, and expand the comprehensiveness of genomic classification. Short-read sequencing (including whole-genome, whole-exome, transcriptome sequencing, or a combination thereof) is an established single assay approach to resolve structural variation associated with B-ALL and AML (Zhang et al., 2016; Liu et al., 2016; Fischer et al., 2015; Paulsson et al., 2015; Leisch et al., 2019). Long-read sequencing approaches offer potential relative advantages, including decreased turn-around time, decreased computational complexity, decreased cost, and improved structural variation detection (Jeck et al., 2019; Liu et al., 2020; Oikonomopoulos et al., 2016; Jain et al., 2018).

Long-read sequencing approaches, like Oxford Nanopore Technologies (ONT) platforms, have the potential to generate faster and more cost-effective results compared with short-read sequencing methods – this is in part due to the ability to generate sequence data continuously and asynchronously such that sequencing starts generating data immediately and can continue until a user-defined limit is reached or until a confident genomic characterization is made.

Adaptive sampling is a nanopore-specific *in silico* enrichment technique that optimizes sequencing by selectively enriching for fusion oncogene targets crucial for understanding disease origins, while maintaining broad genomic coverage necessary for detecting large-scale chromosomal alterations. Adaptive sampling is performed by modulating the current of nanopores during sequencing to keep DNA fragments of interest and physically ejecting fragments not matching a predefined list of targets. The signal generated by each pore is analyzed continuously to identify the source (location) of the DNA in the genome. After one second of sequencing time, the first “chunk” of signal data - 1s of 5Khz signal data corresponding to ∼400nt - is sent to the attached computer. The signal chunk undergoes basecalling and alignment to the reference genome and a decision is made to either keep (if the sequence is near a gene of interest) or eject each read. If an “eject” decision is made, the sequencer is signaled and the voltage bias on the corresponding nanopore is reversed, effectively removing the DNA from the pore and allowing another to enter. In practice, analysis of a chunk and signaling ejection is accomplished in under one second following the second of data collection, resulting in an average ejected read length of ∼600-700nt (1.5 - 2s total). This selective enrichment, resulting in a 10- to 20-fold increase in sequencing depth over enrichment targets, does not rely on targeted sample preparation (ex., biotin probes or selective PCR amplification), and also maintains the sequencing breadth necessary to determine copy-number variation status, and the sensitivity crucial to identifying gene fusions (Weilguny et al., 2023; Martin et al., 2022). ONT sequencing platforms balance focused analysis and comprehensive genomic sequencing, offering a more efficient use of resources and time.

We demonstrate the feasibility of using nanopore-based long-read sequencing as a classification tool in pediatric B-ALL and AML through a combination of retrospective sequencing of clinically validated samples and real-time sequencing of diagnostic samples from UNC Hospitals. We employ a novel bioinformatics pipeline using nanopore adaptive whole-genome sequencing data to infer genetic abnormalities at multiple scales: chromosome-level aneuploidy, large-scale structural variants including inter-chromosomal translocations, and gene-level copy-number variation and small sequence variants. This approach correctly classifies clinically relevant aneuploidy (hyperdiploidy, hypodiploidy), translocations that result in fusion transcripts (ex., *ETV6*::*RUNX1*), as well as complex rearrangements (ex., involving *DUX4* and *IGH*), and subchromosomal copy-number variants (ex., iAMP21, *CDKN2A*, *ERG*, *FLT3*-ITD).

Additionally, we show potential for genotyping of single nucleotide variation (SNV) at pharmacogenomically relevant loci, including *TMPT* and *NUDT15*. In this pipeline, we optimized sample preparation, sequencing, and adaptive sampling parameters to robustly identify fusions and small variants while maintaining the breadth necessary to visualize gross changes in chromosome copy number.

## Materials and Methods

We performed ONT whole genome sequencing (WGS) on fifty-seven (57) acute leukemia specimens representing diverse clinically diagnosed genomic subtypes (retrospective sampling; n=39) or new diagnoses before clinical genomic subtyping (real-time sampling; n=18) (Table 1). DNA from these specimens was extracted from peripheral blood mononuclear cells (PBMCs), bone marrow mononuclear cells (BMMCs), or whole blood (Supplemental Table 1). Samples were obtained from the University of North Carolina at Chapel Hill (UNC) and St. Jude Children’s Research Hospital (SJCRH) with approval by their respective Institutional Review Boards. Illumina RNA sequencing was available for samples from SJCRH. Clinical diagnosis at UNC was determined by G-banding karyotype analysis, FISH, and sometimes microarray as part of the standard of care.

**Table 1.** The number, lineage, and genomic subtype of specimens sequenced in this study using nanopore whole-genome sequencing with adaptive sampling.

| Number of Samples | Lineage<sup>1</sup> | Genomic Subtype |
| --- | --- | --- |
| 9 | B-ALL | High Hyperdiploidy (>50 chromosomes) |
| 4 | B-ALL | Low Hypodiploidy (31-39 chromosomes) |
| 4 | B-ALL | Near-haploidy (24-30 chromosomes) |
| 2 | B-ALL | iAMP21 |
| 6 | B-ALL | *ETV6::RUNX1* |
| 2 | B-ALL | *TCF3::PBX1* |
| 3 | B-ALL | Ph |
| 8 | B-ALL | Ph-like |
| 2 | B-ALL | *MEF2Dr* |
| 3 | B-ALL | *ZNF384r* |
| 3 | B-ALL | *DUX4r* |
| 2 | B-ALL | B-other |
| 1 | AML | *RUNX1::RUNX1T1* |
| 4 | AML | *KMT2Ar* |
| 2 | AML | *NUP98::NSD1* |
| 1 | AML | *FLT3-ITD* |
| 1 | AML | *NPM1*-mutated |

### DNA extraction and shearing

DNA was extracted from cryopreserved or fresh samples using the ZymoBIOMICS MagBead DNA/RNA kit following manufacturer’s instructions (Zymo Research). Fragment sizes in excess of 20Kbp - without significant degradation - were verified by gel electrophoresis. Extracted DNA was sheared using a 26G-1" needle for a total of 7 passes to obtain a fragment size distribution for optimal nanopore sequencing throughput. Size selection was performed using 0.4 volumes of Ampure XP Beads (Beckman Coulter). DNA was quantified using the Qubit fluorometer with the Qubit dsDNA Quantification, High Sensitivity Assay Kit (ThermoFisher Scientific).

### Library preparation and sequencing

Ligation-based library preparation of native DNA was performed using the following Oxford Nanopore Technologies library preparation kits: SQK-LSK 109, SQK-LSK 110, SQK-LSK 112, and SQK-LSK 114, SQK-NBD 114. All library preparation was conducted following the manufacturer’s instructions, with the exception of SQK-LSK 114, which was modified in several ways, including the exclusion of 1uL DNA CS, and a reduction in the final room temperature incubation time of 5 minutes instead of the listed 10 minutes (Supplemental Methods). Samples were sequenced either multiplex or singly on FLO-MIN106, FLO-PRO002, FLO-PRO112, or FLO-PRO114M flow cells for up to 72 hours or until the available pores were exhausted. Samples were sequenced on a PromethION 2 Solo (P2) machine or MinION device – all except four samples were sequenced using adaptive sampling (Supplemental Table 1). Initial sequencing using adaptive sampling enriched for 59 genes frequently involved in B-ALL and AML translocations and fusions (Supplemental Table 2). Subsequently, two larger gene panels were employed: a 152-gene panel was used for six samples and an expanded 223-gene panel for four samples. These enhanced panels were designed to capture additional genes and genetic regions implicated in B-ALL, AML, and T-ALL (Supplemental Tables 3 and 4). To inform the scope of relevant fusions, we referenced previous work detailing the landscape of ALL and AML genomic subtypes (Brady et al., 2022; Umeda et al., 2024) First, single partner genes were parsed from gene fusions detected by RNA-seq or WGS. Genes that occurred as a fusion partner in more than one distinct case were included. We included the entire genomic range for each gene or locus (ex., IGH), as annotated on GRCh38, and a margin of 50Kbp on either side.

Reads were base-called (and de-multiplexed, if applicable) using Dorado (v0.5.1-0.6.0) in super-accurate duplex mode. Base-called reads were aligned to the GRCh38 human reference genome using minimap2 (Li, 2018).

### Digital karyotyping and aneuploidy inference

Relative copy number across the genome at the chromosome level ("digital karyotype") was inferred based on relative sequencing depth by assessing genome-wide and chromosome-level depth of coverage (Figure 1A). Briefly, we infer a baseline diploid (uniform) sequencing depth equivalent to the non-blast percentage (typically low), then assess the relative read depth above baseline, where a 2:3 ratio is observed between diploid and triploid chromosomes and 1:2 between haploid and diploid, respectively. To compare with G-banding karyotypes and assess gross aneuploidy levels, we discuss only whole-chromosome and arm-level gains and losses, although smaller subchromosomal gains and losses are also clearly evident. To avoid the potentially confounding effect of adaptive sampling on relative sequencing depth assessment, we constructed this coarse-scale depth as a function of reads per million base pairs (Mbp), where each read contributes a count of one to the bin in which the center of the read aligns. To assess minimum sensitivity to detect chromosome-level copy number changes, we consider cumulative reads at each timepoint (each minute) after sequencing is initiated – based on read timestamps. We apply a pairwise Kolmogorov-Smirnov test with a conservative threshold (p < 1e-9) and a minimum median divergence of 20% to determine the earliest timepoint at which chromosomes exhibit significantly divergent copy number meeting the clinically-relevant threshold for high hyperdiploid (≥53) or low hypodiploid/near haploid (≤39) karyotype.

**Figure 1.**
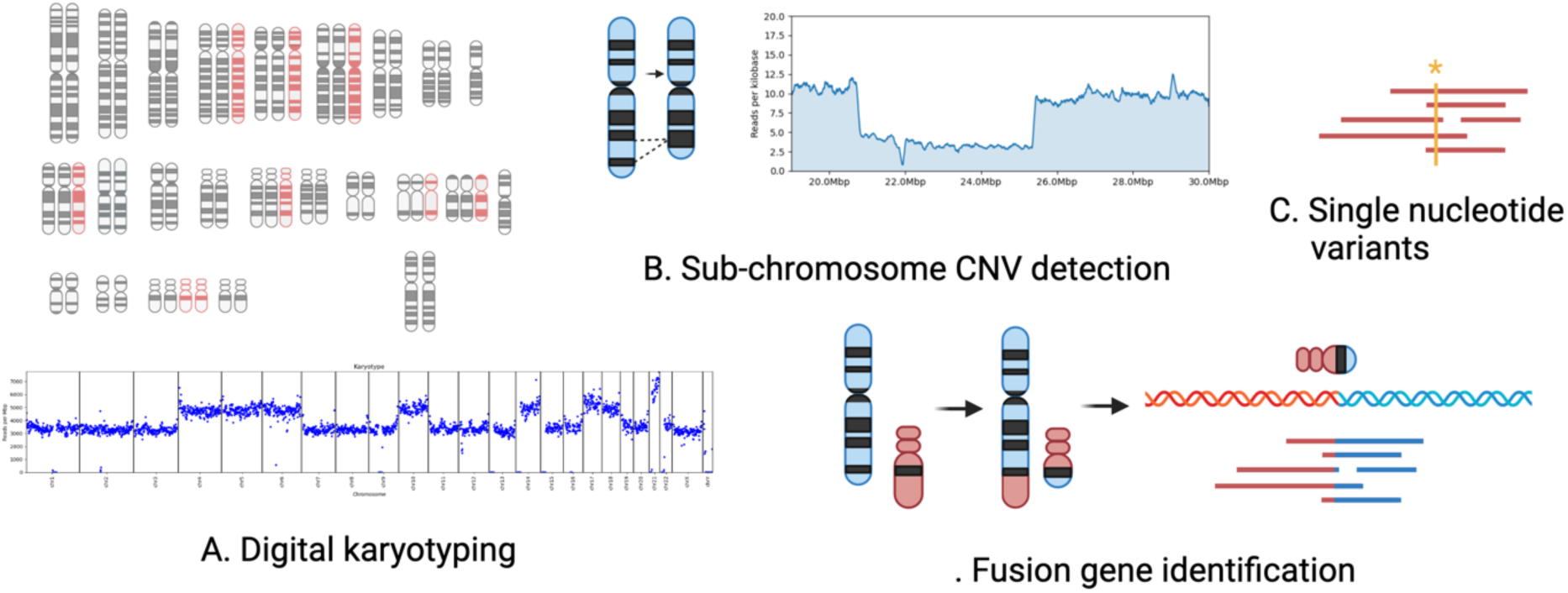
Analytical overview. (A) large-scale copy-number characterization (karyotyping) based on relative chromosome-wide sequencing depth, (B) small-scale/intragenic copy number variation detection based on sequencing depth and split-read mapping, (C) single nucleotide variation (SNV) calling within enriched gene targets, and (D) fusion gene identification based on split alignment of long reads.

### Translocation and fusion detection

Putative translocations were characterized by counting reads for which multiple alignments existed to two independent genes in our enrichment set (ex., *ETV6* and *RUNX1*) (Figure 1D). A read was considered "anchored" in a gene if an alignment of ≥500nt existed within 5Kbp of the gene – this accounted for rearrangements involving the translocation of promoters outside the annotated gene boundaries. Known repetitive elements (Smit et al., 2013-2015), centromere and satellite sequences, and segmental duplications were masked to avoid false positives caused by ambiguous alignments and transposable elements. Translocations between two genes in our target set with at least two independent supporting reads were subsequently validated by visualization of the supporting and putative breakpoints. Fusions detected in our samples had a minimum of 3, and as many as 104 supporting (non-duplex) reads (Supplemental Table 5). Two apparently independent reads can sometimes represent both strands of a "duplex" read that were not appropriately collapsed during duplex basecalling. Duplex reads were identified conservatively and considered a single read if two reads supporting the same translocation were acquired through the same sequencing channel within 30 seconds of one another.

### Small-scale and intragenic copy-number variation

Smaller-scale (sub-chromosomal) copy number variation (ex., iAMP21, *CDKN2A,* and *ERG* deletion, *CRLF2*-*P2RY8* interstitial deletion) was determined by a combination of sequencing depth and split read alignment evidence (Figure 1B). Focal deletions are characterized by a haploid (or multiple thereof) drop in read depth and reads split-mapped to either side of the deletion breakpoints. We consider the full per-nucleotide sequencing depth to evaluate intragenic copy-number variation within genes in our enrichment set.

### Targeted SNV and insertion/deletion calling

Due to uneven coverage resulting from adaptive sampling enrichment and to additionally capture moderate-sized insertions/deletions, existing SNV calling tools for nanopore sequencing data were found to be ineffective. We screened our samples for clinically relevant single nucleotide variants (SNVs), specifically focusing on mutations in *NPM1*, *FLT3*-ITD, and *TPMT*/*NUTD15*. We implemented a straightforward reference-guided assembly approach to identify SNVs and small-scale insertions and deletions (indels) above 0.3 minor allele frequency (MAF). This threshold of 0.3 MAF detected heterozygous and homozygous variants consistent with known molecular genetics across our pediatric leukemia cohort, the majority of which have blast percentages greater than 80%. Briefly, within each enriched gene region, we align all overlapping reads to the GRCh38 reference genome and build a consensus sequence, including SNVs and indels at or above 0.3 MAF. All overlapping reads are subsequently realigned to the consensus sequence and SNV and indel variants exceeding 0.3 MAF are reported. Small-scale indels, notably *FLT3*-ITD, were likewise identified by building a local consensus sequence from a high-depth sequence covering our target genes. We additionally devised a notion of *in silico* PCR to recapitulate commonly used capillary electrophoresis methods to characterize *FLT3*-ITD (Kiyoi et al., 1999) where read segments bounded by *FLT3* 11F (GCAATTTAGGTATGAAAGCCAGC) and 12R (CTTTCAGCATTTTGACGGCAACC) primers were extracted and plotted by size (Figure S1).

### Evaluation and validation

Leukemia lineage was determined by flow cytometry. The ground truth of subtypes was determined by G-banding karyotype analysis, FISH, microarray, Illumina sequencing, or some combination thereof. Illumina RNA sequencing and fusion detection and/or expression-based subtyping (ex., *DUX4*r) determined a subset of cases. We evaluated the performance of our WGS-based analysis against the final consensus subtype following this multimodal characterization.

### Cost analysis

To evaluate the cost of adaptive nanopore sequencing, we conducted a microcosting analysis of reagents and consumables. Based on the sequencing depth, we conducted a costing sensitivity analysis to account for uncertainties associated with practical implementation. Activity-based costing for human resources was excluded as sunken costs for the purposes of this analysis.

### Code availability

Software used for analysis is publicly available at https://github.com/jwanglab/long-read-leukemia-dna.

## Results

Nanopore-based whole genome sequencing demonstrated 100% specificity and 96% sensitivity in aggregate for determining genomic subtype across both retrospective and real-time samples. The choice of library preparation method and sequencing platform affected assay performance only through their influence on read length and sequencing coverage. Of the 57 samples analyzed, we accurately characterized gross karyotype abnormalities in all but two cases, achieving 96% sensitivity. Both occurred in specimens with very low (<30%) blast content; among specimens with >50% blasts (53/57), we achieved 100% accuracy. Our approach achieved 100% specificity in the sense that no gross aneuploidy was reported for cytogenetically diploid cases.

Nanopore sequencing successfully identified all fusion events previously detected by clinical testing, achieving 100% sensitivity for detection of known fusion oncogenes. Moreover, in five cases, this approach revealed additional fusion partners that were not identified through clinical karyotyping and FISH. These included cytogenetically cryptic (ex. *DUX4*::*IGH*) fusions and rearrangements where only one gene partner was known (ex. *PDGFRB*/*CSF1R* rearranged) that were fully resolved by nanopore sequencing. In all cases, nanopore-based translocations were consistent with available clinical cytogenetic evidence, achieving 100% specificity.

Although genome-wide SNV calling was not performed in clinical testing, our sequencing revealed the correct SNV in each of the 4 cases with a clinically identified SNV - clinically relevant SNVs including *TMPT* and *NUDT15* were assessed for in each nanopore-generated dataset regardless of positive clinical findings (Supplemental Table 6). Across all cases, nanopore WGS with adaptive sampling confirmed all but two clinically identified genomic alterations used for risk stratification and treatment planning. Additionally, this method revealed *DUX4* rearrangements in two cases that were not detected by standard clinical testing, potentially warranting revised risk assessments for these patients.

For samples run singleplex with adaptive sampling on a P2, we generated an average of 12X whole-genome coverage and 86X coverage over target genes. For all samples run with adaptive sampling (including multiplexed aneuploid samples), we achieved a relative enrichment of 8.3X (range 1.46 - 16.4X). Samples exhibiting very poor enrichment resulted from severely fragmented input DNA (0154, 0160) or adaptive sampling failure (0229). Samples undergoing adaptive sampling from fresh samples for real-time analysis ranged from 4.2 - 12.5X enrichment.

### Nanopore whole genome sequencing with and without adaptive sampling accurately identifies clinically relevant karyotype profiles and gene fusions in acute leukemias

The karyotype for 27 of 57 samples showed gross changes at the chromosome level (aneuploidy) (Figure 2). Clinically relevant subtypes classified included high hyperdiploidy (>50 chromosomes; n=9), low hypodiploidy (31-39 chromosomes; n=4), and near-haploidy (24-30 chromosomes; n=4) (Table 1). Ten additional samples were aneuploid (45-50 chromosomes) with (n=8) or without (n=2) other known genomic drivers (Supplemental Table 1). In all 27 aneuploid cases, changes in gross karyotype detected by WGS nanopore sequencing (both with (n=25) and without (n=2) adaptive sampling) were consistent with clinical classification. 2/57 samples had low blast count (<30%) that prohibited precise genomic classification (samples 0131, 0133) (Figure S2). In cases 0154 and 0164, we estimated six copies of *RUNX1* within a broader regional amplification pattern (Figure S3), indicating an intrachromosomal amplification of chromosome 21 (iAMP21) B-ALL genomic subtype, defined by at least four copies of *RUNX1* and focal amplification.

**Figure 2.**
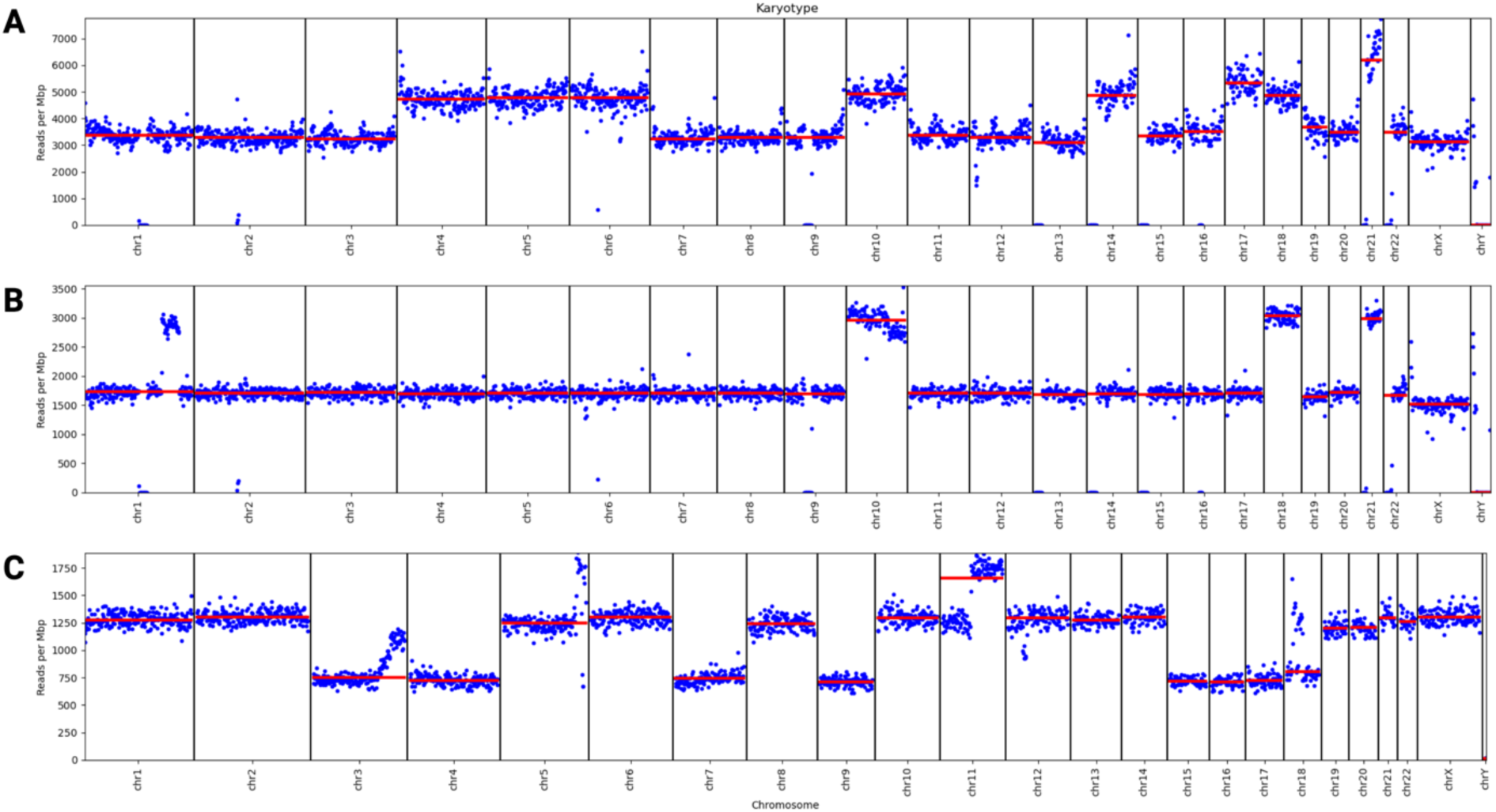
Digital karyotypes are inferred based on the relative number of reads per chromosome; blue points show the number of reads in non-overlapping 1Mbp bins and red lines indicate the median for the entire chromosome. (A) Patient sample 0052 represents a case of B-ALL with a high hyperdiploid genomic subtype. Our inferred digital karyotype is 55, XY, +4, +5, +6, +10, +14, +17, +18, +21, +21. (B) Sample 0225 represents B-ALL with a near haploid genomic subtype. Our inferred digital karyotype is 27, XY, +10, +18, +21. (C) Sample 0223 is B-ALL with low hypodiploidy, inferred to be 38, XX, -3, -4, -7, -9, +11q, -15, -16, -17, -18. In all cases, we only annotate whole-chromosome and arm-level changes for the purposes of gross aneuploidy detection, but smaller sub-chromosomal copy-number changes are also conspicuous.

We detected 19 unique gene fusions across all samples (Supplemental Table 1; Supplemental Table 5). Identified fusion-driven genomic subtypes of B-ALL were: *ETV6*::*RUNX1*, *TCF3::PBX1*, Philadelphia (Ph) (*BCR*::*ALB1)*, Ph-like including *CRLF2*r (*CRLF2::IGH; PAX5::MLLT3; MEF2D::CSF1R; EBF1::PDGFRB*), MEF2Dr (*MEF2D*::BCL9; MEF2D::HNRNPUL1), ZNF384r (*ZNF384::EP300; TCF3::ZNF384; CREBBP::ZNF384*) and DUX4r (*DUX4::IGH*). Fusion-driven genomic subtypes of AML identified (n=3) were: *RUNX1::RUNX1T1*, KMT2Ar (*KMT2A::MLLT1, KMT2A::MLLT10; KMT2A:USP2*), and *NUP98::NSD1* (Table 1; Supplemental Table 1). Gene fusions within B-ALL fell into two major categories: balanced translocations and complex rearrangements (ex., *DUX4*r; *CRLF2*r). The primary characterization of structural rearrangements from long-read WGS consists of reads aligning to distant genes or regions. Adaptive sampling enriches for reads spanning involved genes, providing strong and consistent support for these structural variants (Figure 3; Figure 4). In several instances, nanopore-generated WGS data provided additional information about structural variation that was not detected through clinical assays. In one case, clinical classification identified one of two fusion partners, whereas nanopore WGS identified both fusion partners – for sample 0130, clinical testing classified sample 0130 as *PDGFRB* or *CSF1R* with an unknown fusion partner while sequencing data identified a precise *MEF2D*::*PDGFRB* fusion. In other cases, clinical break-apart FISH assays only identified one component of the likely fusion (*NUP98* in sample 0158 and *KMT2A* in sample 0172), while the partner gene with prognostic significance was only identified after nanopore sequencing, (*NUP98*::*NSD1* and *KMT2A::USP2*). In two cases (0157 and 0162), an *IGH*::*DUX4* fusion (which is karyotypically cryptic) was not reported clinically (by karyotyping and FISH) but was detected in our real-time sequencing analysis.

**Figure 3.**
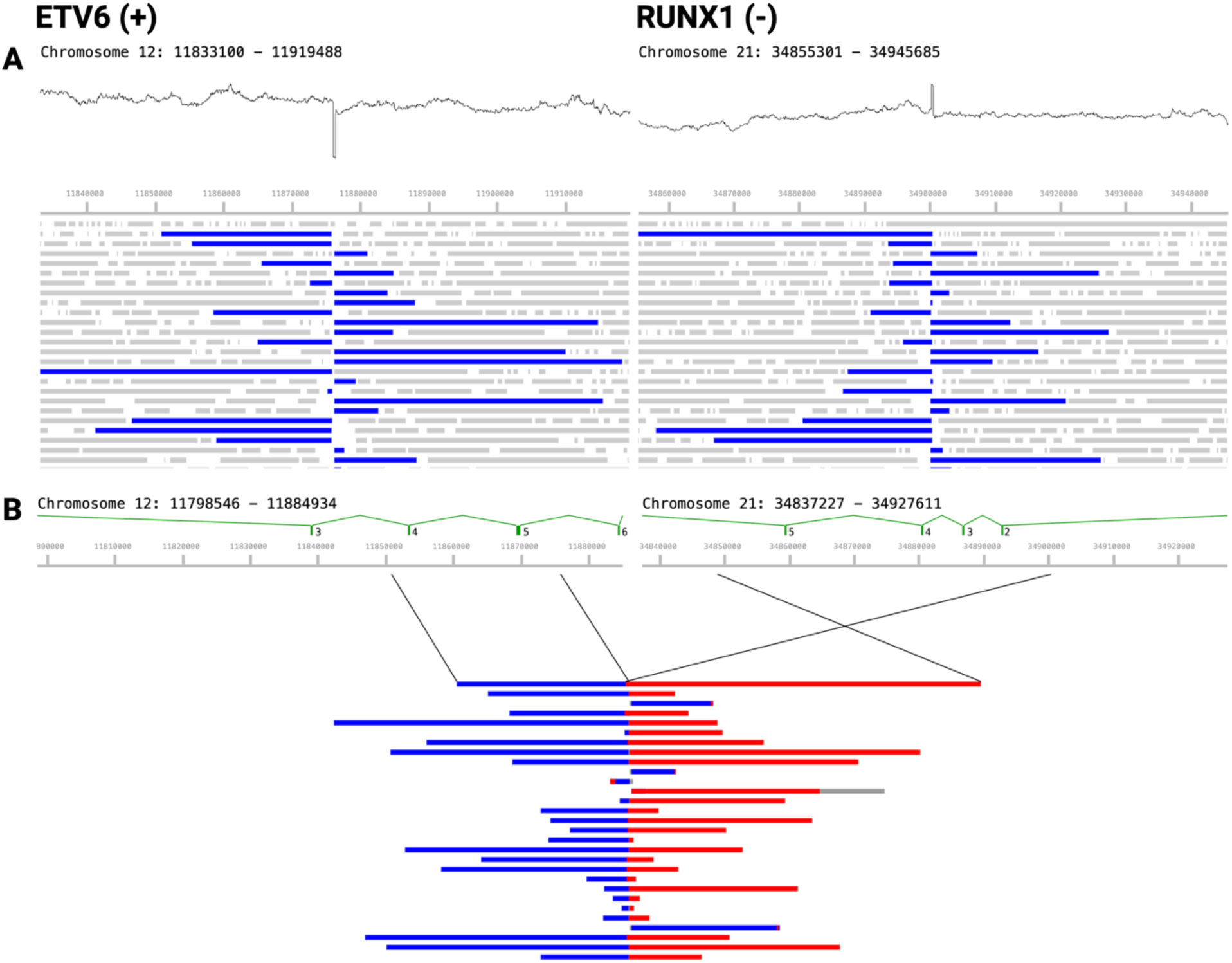
Sample 0250 represents a case of B-ALL with an *ETV6*::*RUNX1* genomic subtype. (A) Long reads aligning to *ETV6* (left) and *RUNX1* (right); blue bars represent matched fragments overlapping the putative breakpoint; gray bars are singly-mapping. Relative sequencing depth is also shown above, indicating discontinuous coverage representing small indels at the putative translocation breakpoint resulting from imperfect double-strand break repair. (B) A sample of reads supporting the putative *ETV6*::*RUNX1* breakpoint, showing segments mapping to *ETV6* (blue) and *RUNX1* (red) indicating an inverted translocation consistent with *ETV6* intron 5-6 fused to *RUNX1* intron 1-2, matching their respective coding orientation.

**Figure 4.**
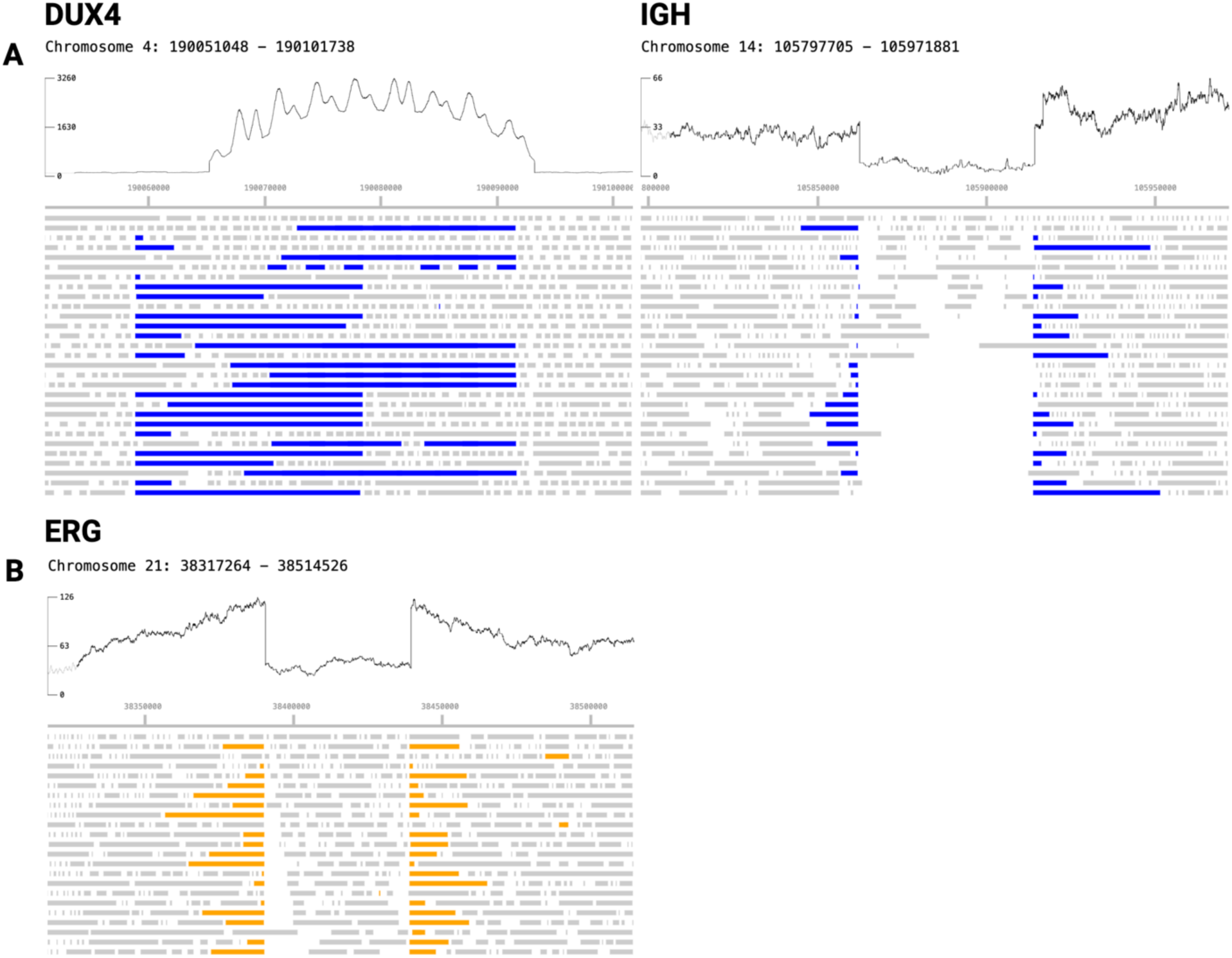
Sample 0162 with *DUX4*-*IGH* rearrangement (A) where blue bars indicate reads aligning to both regions, (B) with a focal deletion in *ERG* characterized by a drop in sequencing depth and reads split across the deletion boundaries (orange).

### Small-scale and intragenic copy-number variation and single nucleotide variation are sensitively characterized in enriched regions

Nanopore WGS produces support for clinically relevant small-scale structural variants affecting genes associated with ALL and AML, including *CRLF2*-*P2RY8* interstitial deletion (0060, 0136, 0163, 0238), partial or total loss of *CDKN2A* (0153, 0165), and heterogeneous *ERG* deletion (0162). A full list of observed small variants, including CNVs and SNVs is provided in Supplemental Table 6. We accurately detect these deletions with a combination of sequencing depth and long reads spanning the deletion boundaries (Figure 4B).

We assessed the utility of adaptive WGS to call single-nucleotide polymorphisms (SNPs) and small insertions by examining pharmacogenomically relevant SNVs in *TPMT*, and *FLT3* internal tandem duplication (*FLT3*-ITD) - a driving mutation in pediatric AML (Table S5). We identified relevant SNVs in *TPMT* in both cases (0136, 0162) with clinically identified mutations (A154T, Y240C). These are trivially phased in our long reads and confirmed to occur on the same haplotype. *FLT3*-ITD was called in one AML case (0141), characterized by a consensus tandem duplication of *FLT3* CDS loci 1823-1904 (81nt). To recapitulate commonly used PCR and capillary electrophoresis detection of *FLT3*-ITD (Kiyoi et al., 1999), we identified 46 reads that include *FLT3* 11F and 12R primer sites. The size distribution of these ITD-spanning reads is shown in Figure S1, producing an ITD:WT allelic ratio (AR) of 0.64, consistent with the clinically reported AR of 0.65.

### Clinically relevant genomic variation and tumorigenic drivers are robustly detectable with a single rapid, low-cost assay

Real-time sequencing and analysis generates sample classifications within 9 hours of sample receipt. DNA extraction and shearing using the ZymoBIOMICS MagBead DNA/RNA kit took less than 2 hours, and library preparation took approximately 75 minutes. Using the high-throughput PromethION flow cells, a digital karyotype could be inferred within 15-30 minutes of the start of sequencing. Fusion detection (defined as two or more independent reads supporting a single given fusion) typically took between 3 and 6 hours from the start of sequencing (Fig. 5). In concordance with our retrospective sampling, real-time classification using our nanopore-based WGS approach was 100% consistent with clinically-derived genomic subtype classification.

**Figure 5.**
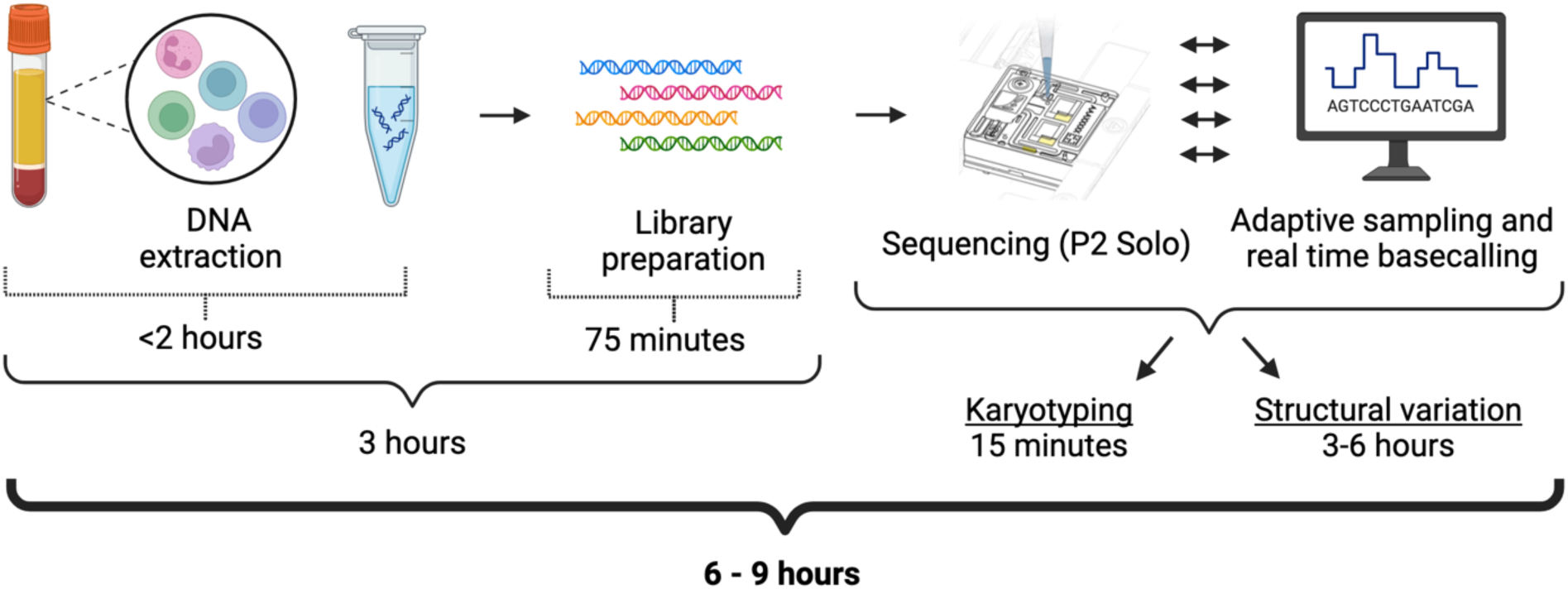
Real-time sample processing and analysis timeline. DNA extraction and library preparation took 3 hours, and sample classification took between 3 and 6 hours. Complete sample classification (from sample receipt to classification) took between 6 and 9 hours.

To validate detection limits, we analyzed three sample sets multiplexed on individual flow cells: two sets of seven samples with known aneuploidies and one set of four samples with known translocations. For the aneuploid and fusion-positive samples, we obtained an average of 8.6X and 23.6X coverage of enriched targets, respectively.

These are broadly as expected given the level of multiplexing. In all cases, we successfully identified the clinically determined genomic subtype. We retroactively assessed the timing of translocation detection across the entire cohort. Briefly, since nanopore sequencing reads are captured asynchronously and continuously from the time sequencing is initiated, and each read is time-stamped with the time it started and finished sequencing, we can infer the total duration of sequencing necessary to call a translocation based on the time the second independent supporting read finished sequencing. Analysis of sequencing data is performed continuously in real time and adds a negligible time for analysis after the second read is complete - we conservatively add five minutes. Figure 6 shows the distribution of minimum necessary sequencing times to conclusively identify translocations using this method. When using adaptive sampling and running samples individually, 19/25 (76%) of translocations are identified in under 3 hours, and 23/25 (92%) are identified in under 6 hours (Fig. 6A). As expected, samples multiplexed on one flow cell or run without adaptive sampling take longer to detect the translocations. Two samples that were sequenced individually with adaptive sampling are apparent outliers, taking 11-14 hours to result. Sample 0136 is one, taking 11.2 hours to identify the putative driving translocation (*CRLF2*::*P2RY8*) based on the clinical diagnosis, however we also find a second putative driver, *PAX5*::*MLLT3*, in only 2 hours. For the other, Sample 0231, we detected *MEF2D*::*BCL9* in 13.9 hours; the total sequencing efficiency and throughput was lower than typical for this sample resulting in the longer detection time.

**Figure 6.**
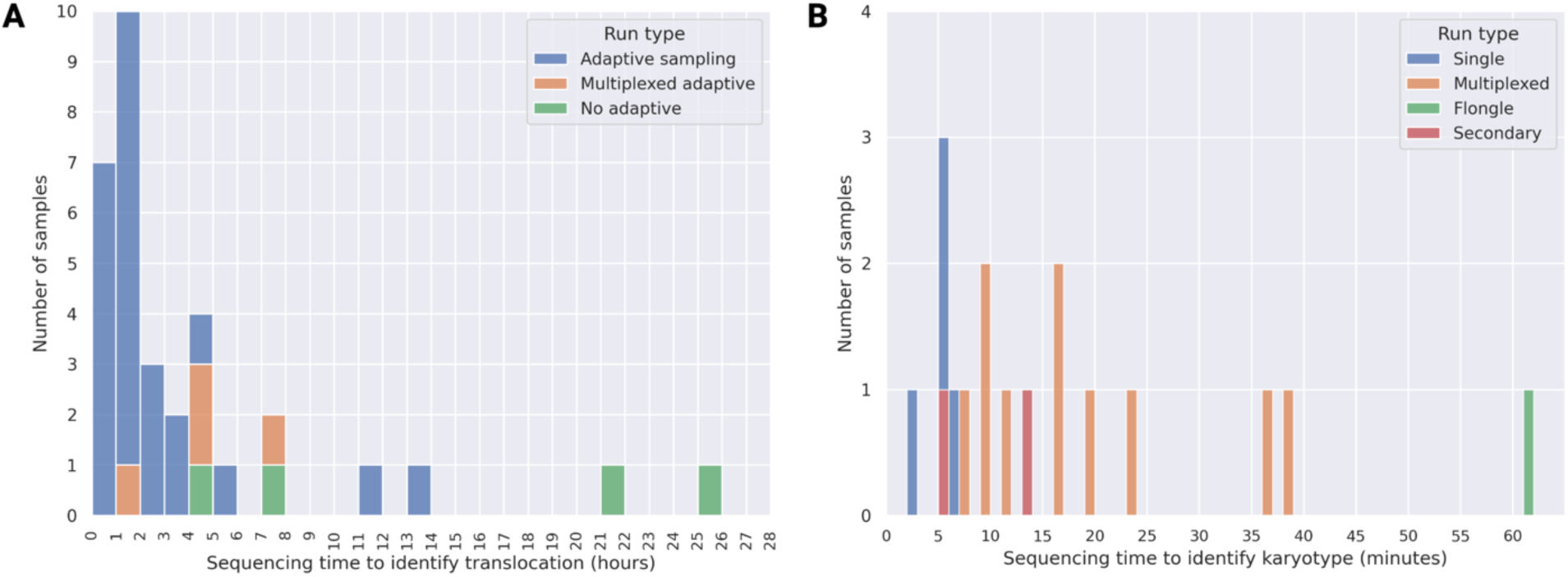
Distribution of total sequencing time required to identify translocations (A) and karyotype (B). (A) When using adaptive sampling and running samples individually (blue bars), 19/25 (76%) of translocations are identified in under 3 hours, and 23/25 (92%) are identified in under 6 hours. Samples run multiplexed (4 on one chip; orange) or without adaptive sampling (green) show the expected longer sequencing times to result. (B) Sequencing time required to classify a chromosome abnormality into a specific aneuploid category (such as high hyperdiploid, low hypodiploid, or near haploid). Blue bars represent samples that were sequenced individually on a P2, orange bars indicate samples that were multiplexed, and one sample (green) was sequenced on a low-throughput Flongle flow cell. Red bars indicate samples sequenced individually where karyotype abnormalities represented subclones secondary to a translocation-driven primary clone (*BCR*::*ABL1*).

We additionally determined the minimum amount of time needed to accurately detect and characterize chromosomal abnormalities by analyzing statistically and clinically relevant changes in chromosome copy number. We excluded the two specimens with very low blast count (<30%), as previously described. Among cases with gross karyotype abnormalities as primary genomic subtypes (high hyperdiploid, low hypodiploid, near haploid), four were sequenced individually on a high-capacity flow cell, one individually on a low-capacity Flongle flow cell, and the remainder multiplexed on a P2 flow cell. Singleplex high-throughput runs were characterized into aneuploidy classes after a median of 5 minutes and a maximum of 6 minutes (Fig. 6B). Multiplexed specimens were called after a median of 16 and maximum of 39 minutes. The specimen run on a low-throughput Flongle flow cell (0048) was called after 61 minutes. Illustrative sequencing depth profiles at this minimum detection timepoint are shown in Figure S4. Real-time analysis and assessment of karyotype abnormalities showed that these signatures are robustly detected within minutes of the start of sequencing. However, in all cases sequencing continued well past this point to identify possible concomitant translocations (ex. *BCR*::*ABL1*).

We performed a microcosting exercise to provide information on the costs associated with this proposed single assay approach. Our evaluation encompassed the expenses associated with consumables and reagents for both extraction and library preparation. We also factored in the cost of flow cells, estimated based on a moderate bulk purchase of 32 units. Given that the required sequencing depth may vary across genomic subtypes, we performed a sensitivity analysis to account for this uncertainty. Our findings revealed that when dedicating an entire flow cell to a single specimen, the material costs amounted to $934.64. Based on computational downsampling analysis and testing of multiplexed specimens, we demonstrated that up to four specimens can be effectively sequenced on a single flow cell while retaining high sensitivity.

Specifically, the material cost per specimen decreased to $483.18 when processing two specimens per flow cell, and further reduced to $249.06 when running four specimens per flow cell (detailed breakdown available in Supplemental Table 7). While further validation is needed, this cost structure demonstrates the potential for significant economies of scale, offering flexibility in balancing cost-efficiency with the specific requirements of genomic analyses.

## Discussion

Advancements in genomic classification of acute leukemia over the past decades have led to improved prognostic stratification, risk-adapted therapeutic selection, and precision therapy. While some treatment centers perform comprehensive genomic profiling of all patients with acute leukemia, this is the exception. At most treatment centers in high-income countries (HICs), genomic classification continues to rely on long-established techniques in cytogenetics, supported by targeted molecular profiling. More importantly, most cancer treatment centers globally are in low and middle-income countries (LMICs), often without access to reliable cytogenetic testing approaches. Therefore, accessible, scalable approaches to improve clinical genomic classification of acute leukemia are critically needed.

The motivation to pursue nanopore adaptive WGS is for simplified identification of well-established clinically useful genomic results, as opposed to a focus on discovering new biological insights. The results presented suggest that, as a single assay classification tool, nanopore-based adaptive whole-genome sequencing accurately classifies B-ALL into genomic subtypes, with the potential to identify clinically relevant AML genomic subtypes, as well as clinically actionable pharmacogenetic subtypes. We have demonstrated proof of principle that nanopore long-read whole genome sequencing can provide all clinically relevant genomic information currently offered by traditional diagnostic testing (karyotype, FISH, and occasional microarray) for pediatric acute leukemia. In our cohort, there was no loss in sensitivity or throughput with a variety of sample preparations – robust results were generated from freshly collected as well as cryopreserved samples, including whole blood, isolated mononuclear cells, bone marrow aspirate, and peripheral blood (Supplemental Table 1). Likewise, nanopore-based WGS is less reliant on high-quality viable cells that are beneficial for karyotype and FISH analysis. Our approach comprehensively identifies known variation commonly characterized by a combination of karyotyping, FISH, and targeted molecular tests.

### Capturing complex and cytogenetically cryptic rearrangements

The long reads produced by nanopore sequencing are particularly useful for identifying complex structural rearrangements that are difficult to identify with current clinical approaches, such as *DUX4* rearrangements. *DUX4* rearrangements make up about 14% of B-ALL cases (Lee et al., 2021), yet structural variations involving *DUX4* are often not well characterized due to tandem *D4Z4* repeat cassettes containing *DUX4* (Rehn et al., 2020). The identification of *DUX4* rearrangements commonly relies on the detection of fusion transcripts by RNA sequencing (Bařinka et al., 2022). We detected *DUX4*::*IGH* gene fusions in three samples (one retrospective sample, 0222, and two real-time samples, 0157 and 0162) that were not identified by conventional FISH or karyotype analysis. Under standard clinical classification, these cases would have been designated as risk-neutral. However, emerging evidence suggests that *DUX4* fusions may indicate a more favorable prognosis. These findings demonstrate that nanopore adaptive WGS can reveal clinically relevant genomic alterations even in settings with sophisticated diagnostic capabilities.

### Small variant characterization

This assay provides a level of characterization of aneuploid and small copy number changes that is critical for clinical decision-making. While hyperdiploid karyotype is generally a favorable prognostic factor, multiple groups have demonstrated that the prognosis is influenced by specific chromosome gains. More recently, multiple groups have demonstrated the potential prognostic value of small deletions in certain contexts, specifically *IKZF1* deletion (Boer et al., 2016; Mullighan et al., 2009). It is important for future genomic classification approaches to B-ALL to include small deletions in the diagnostic results.

Pharmacogenomics is a rapidly expanding field with increased clinical relevance. For patients with ALL, the key pharmacogenomic information needed is *TPMT* and *NUDT15* genotype, which has direct clinical implications. In HICs, the genotype is typically determined by a targeted molecular assay. We show the potential for Nanopore adaptive WGS to detect known SNPs in *TPMT* with the same assay that provides genomic classification of B-ALL. This provides another example of cost savings in HIC and the expansion of clinically relevant pharmacogenomic information in LMICs.

### Digital karyotyping performance and limitations

Nanopore-based WGS is a promising approach for the classification of B-ALL, yet accurate classification is constrained in cases with low blast percentages and is sometimes limited to subtypes defined by genomic structural variation. For instance, gains of chromosomes 4, 6, 14, 17, 18, and 21 were detected by microarray (but not traditional karyotyping) in sample 0133 – these gains were largely undetectable with our current analytical pipeline. Likewise, with patient sample 0131 (near haploid), gains of chromosomes 21 and X were shown through clinical testing but were undetectable in nanopore WGS (Supplemental Figure 3). Additionally, a small portion of acute leukemia cases are defined by expression profiles without subtype defining DNA variation, such as the proposed *ETV6*-like, *KMT2A*-like, and *ZNF384*-like cases, limiting the utility of DNA-based classification approaches in these cases (Gu et al., 2019).

There are, in several cases, large-scale structural variants – including whole-chromosome gains and losses – that are detected by our nanopore sequencing that are inconsistent with the reported clinical G-band karyotype (Table S1). Because clinical karyotyping involves growing cells in culture, genetic drift may occur, meaning that both the original (G-band karyotype) and observed (nanopore-derived) genetic classifications could be valid representations of the cells at different points in time. Previous digital karyotyping approaches based on genomic data have noted similar discrepancies in a minority of cases, emphasizing possible G-band karyotyping inaccuracies (Bařinka et al., 2022). However, these differences are typically minor and do not result in a different clinically-relevant aneuploidy class (high hyperdiploid, low hypodiploid, near haploid).

### Implementation and future directions

Iterative adjustment of this sequencing approach to focus on B-ALL, all acute leukemia, or solid malignancies, would be made without modifications to wet lab sample preparation. The differences in genomic targets and data analysis required only minimal informatic changes. The simplicity in reagents and wet lab training across focused assays offers a large advantage in the supply chain, human resources, cost, and speed of assay improvements. The material costs, which we provided in a sensitivity analysis across assay throughput and depth, compares favorably with the traditional combined multiple technology approach of karyotype, FISH, targeted molecular approaches traditionally used to determine genomic subtypes of acute leukemia.

Optimization and automation of the sequencing pipeline are essential to democratizing this approach, thereby increasing throughput and decreasing the material and labor costs associated with classification. We will optimize the assay by further exploring the limits of detection based on both blast percentage and sequencing depth. The sequencing depth required is crucial to optimizing cost and throughput with particular implications for lower-resourced settings. Currently, karyotyping and determination of small deletions or gains using nanopore-based WGS requires inferring a karyotype based on the relative number of reads per chromosome. Future development will focus on expanding our automated capabilities beyond fusion calling, particularly for karyotype analysis and indel detection. Translation to clinical practice will involve validation studies in a CLIA-certified laboratory.

## Acknowledgments

This work was supported by the National Institutes of Health R21CA259926 to TBA, R01CA293366 to JRW, and support from the UNC Lineberger Comprehensive Cancer Center and the University Cancer Research Fund.

## Author Contributions

JRW, NB, and TBA conceived and designed the study. JG, KO, and JRW performed sequencing experiments. LR, CM, and TBA acquired specimens and clinical data. JG, JL, and JRW performed analysis and evaluation. JG, NB, TBA, and JRW wrote and edited the manuscript. All authors provided feedback and edited the manuscript.

## Data Availability Statement

Supplemental tables 1-6 include specimen and sequencing characteristics, enrichment gene set, characterized translocations, copy-number variants, and SNVs across this cohort. Sequencing data is available from the authors on reasonable request.

