## Supplemental material for "Real-time genomic characterization of pediatric acute leukemia using adaptive sampling"

**Supplemental Tables:**

**Table S1.** Sample metadata including clinical genomics, sequencing characteristics, and sequencing-based copy-number and fusion drivers.

**Table S2.** List and coordinates of genes used in 59-gene set enrichment

**Table S3.** List and coordinates of genes used in 152-gene set enrichment

**Table S4.** List and coordinates of genes used in 223-gene set enrichment

**Table S5.** Translocation/fusion genes and read-level support

**Table S6.** Focal copy-number variation and SNVs in enriched clinically relevant gene targets

**Table S7.** Microcosting analysis

**Supplemental Methods:**

*Protocol - Leukemia Nanopore WGS with Adaptive Sampling*

**Materials**

Kits and reagents

- ZymoBIOMICS MagBead DNA/RNA Kit ([R2136](https://www.zymoresearch.com/collections/zymobiomics-dna-rna-kits/products/zymobiomics-magbead-dna-rna))
- Oxford Nanopore Technologies Ligation Sequencing Kit V14 ([SQK-LSK114](https://store.nanoporetech.com/us/ligation-sequencing-kit-v14.html)) (store at -20°C)
- Flow Cell Priming Kit [(EXP-FLP004)](https://store.nanoporetech.com/us/flow-cell-priming-kit-004.html) (store at -20°C)
- [AMPure XP Beads](https://www.beckman.com/reagents/genomic/cleanup-and-size-selection/pcr) (store at 4°C)
- Qubit dsDNA High Sensitivity Quantification kit ([Q32851 or Q32854](https://www.thermofisher.com/order/catalog/product/Q32854)) (store at 4°C)

Consumables

- PromethION Flow Cell ([R10.4.1](https://store.nanoporetech.com/us/promethion-flow-cell-packs-r10-4-1-m-version.html)) (store at 4°C)
- 100% EtOH (molecular biology grade/nuclease-free) (store at RT)
- Nuclease-free water (NFW) (store at RT)
- 1.5mL microcentrifuge tubes, nuclease-free (recommended: [Eppendorf DNA Lo-bind](https://www.eppendorf.com/us-en/eShop-Products/Laboratory-Consumables/Tubes/DNA-LoBind-Tubes-p-022431021))
- 2mL microcentrifuge tubes, nuclease-free (recommended: [Eppendorf DNA Lo-bind](https://www.eppendorf.com/us-en/eShop-Products/Laboratory-Consumables/Tubes/DNA-LoBind-Tubes-p-022431048))
- Filtered, sterile tips capable of pipetting the full range of 1-1000⎧L
- 3mL syringe ([must be compatible](https://www.bd.com/en-eu/offerings/capabilities/syringes-and-needles/injection-syringes/bd-plastipak-3-piece-syringe) with 26G needles)
- [26G-1”](https://www.amazon.com/26Ga-inches-Individually-Packaged-Pack/dp/B0C33FTYS3/ref=sr_1_3?keywords=26+gauge+needle&qid=1706537844&sr=8-3) needles

Equipment

- [Magnetic rack](https://biologixusa.com/products/Super-Magnetic-Stand-16-Well-with-Locker-Silver-1-Piece-Pack-p371526951?utm_source=googleshopping&utm_medium=shp&utm_network=x&utm_mobile=0&utm_creative=&utm_position=&utm_random=17710823553907987777&gclid=CjwKCAiAtt2tBhBDEiwALZuhAGHPobevNRzSyoMQDLzsQe00nWMaAxFr7wrj6qQNcAAXT8csGFY3zhoC9bgQAvD_BwE&utm_campaign=smart%20shopping%20-%20us%20(1)&utm_ad_group_id=513113&utm_campaign_id=646822&utm_prod_id=371526951&gad_source=1) for 1.5mL tubes
- Heat block for incubations (example: [digital heating drybath](https://www.thermofisher.com/order/catalog/product/88880030))
- PromethION 2 Solo sequencer
- Computer workstation:
  - Please see the recommended computer (and GPU) specifications for the P2: <https://community.nanoporetech.com/requirements_documents/promethion-2s-it-req.pdf>
  - We recommend the following minimum specifications:
    - 8TB+ internal SSD
    - NVIDIA A6000 GPU or better (we use NVIDIA RTX 3090 with 24GB RAM and it works well for adaptive sampling)
    - 64+ GB DDR4+ RAM
    - 12+ core/24+ thread Intel i7/i9 or AMD Ryzen processor
- Qubit or Quantus [fluorometer](https://www.fishersci.com/shop/products/quantus-fluorometer/PRE6150?srsltid=AfmBOornF8mYrefvpDo_scsJLnNwfAqmcbLzP_-72SlnDEsvAn_LxSqsA_g)
- Hula mixer (or comparable gentle rotating mixer)
- Vortexer
- Pipettes capable of pipetting the full range of 1-1000⎧L
- Dual rotor personal microcentrifuge (example: [#2641-0016](https://www.usascientific.com/dual-rotor-personal-microcentrifuge/p/2641-0016))
- -80°C freezer
- 4°C refrigerator

**Protocol**

DNA extraction

Use the [**ZymoBIOMICS Magbead DNA/RNA Kit**](https://www.zymoresearch.com/collections/zymobiomics-dna-rna-kits/products/zymobiomics-magbead-dna-rna), following the manufacturer’s instructions.

DNA shearing & bead wash

1. Make the extracted DNA volume up to at least 200μL
2. Draw the sample up through the syringe with a 26G needle and gently press down on the plunger to eject the liquid back into the 1.5mL tube. This equals one “pass”
3. Repeat this process 6 more times, for a total of 7 “passes”
4. After shearing the DNA, pull apart the syringe and pipette any remaining liquid left in the plastic. This helps to recover some of the liquid that will be lost during the shearing process
5. Add 0.4X well-mixed AMPure XP beads to the sheared sample (if 175μL was retained, add 70μL AMPure XP beads) and mix well on a gentle rotator (or hula mixer) for 5 minutes
6. Briefly spin down the sample and pellet the beads on a magnet until the supernatant is clear and colorless (this should take about 1 minute)
7. Pipette off the supernatant and discard
8. Add 200μL fresh 70-80% ethanol and rotate the tube side to side in the magnet stand to make sure the solution washes over the beads
9. Pipette off the ethanol, ensuring that the beads remain pelleted, and discard
10. Repeat steps 7 and 8
11. Spin down the tube and place the tube back on the magnet. Pipette off any residual ethanol and allow it to dry for about 30 seconds, but not to the point of cracking. If the pellet gets too dry, the DNA will have trouble eluting off the beads
12. Elute into 49μL NFW

DNA quality control

1. Add 199μl 1x dsDNA HS working solution to new 0.5mL tube
2. Add 1μl of sheared, cleaned DNA
3. Mix DNA + dsDNA HS solution briefly by vortexing
4. Read on Qubit (dsDNA protocol)

*The sample should contain at least 600ng (total) of DNA—this is the minimum required to begin library preparation. As an additional QC step, the sample can also be run on a gel to verify DNA size.*

Library preparation

*For single samples, use Oxford Nanopore Technologies’* [*SQK-LSK114*](https://community.nanoporetech.com/docs/prepare/library_prep_protocols/genomic-dna-by-ligation-sqk-lsk114/v/gde_9161_v114_revu_29jun2022) *protocol with the following modifications. Batched samples can be prepared using* [*SQK-NBD114-24*](https://community.nanoporetech.com/docs/prepare/library_prep_protocols/ligation-sequencing-amplicons-native-barcoding-v14-sqk-nbd114-24/v/nba_9168_v114_revl_15sep2022)*, with no modifications.*

Modifications & notes:

**DNA repair and end prep**

- All library preparations began with between 616 and 1,232ng of extracted, sheared, and bead-cleaned DNA, which assumed an average fragment length of 10kb.
- DCS was not used – DCS volume was made up with NFW

**Adapter ligation and clean-up**

- Use 200μL LFB instead of 250μL LFB
- Elute samples in EB for 5 minutes at RT

**Priming and loading the flow cell**

*No modifications*

**Sequencing**

*Use the following sequencing parameters:*

**Run options**

Run limit: 72 hours

Minimum read length: 20bp

Adaptive sampling: GRCh38.fa

BED file: regions.bed (see Supplemental Table 4)

**Analysis**

Basecalling: On

Modified basecalling: Off

Barcoding: Off

**Output**

Location: Choose an SSD with a minimum of 2TB free space – a very high-performing run may produce up to 2TB raw data

Raw reads: On (.POD5, 4000 reads per file)

*Adaptive sampling, which enriches for targeted loci, can be done by inputting the parameters in blue.*

Post-sequencing

The used flow should be washed using [EXP-WSH004](https://store.nanoporetech.com/us/flow-cell-wash.html) and stored at 4°C.

**Supplemental Figures:**

**Figure S1.** Recapitulating commonly used *FLT3*-ITD PCR and capillary electrophoretic analysis using *in silico* primer capture—sample 0141 with clinically reported AR 0.65. We identified 46 reads spanning *FLT3* primers, 28 consisting of 300-309nt and 18 containing 377-392nt, inferring an ITD:WT ratio of 0.64.

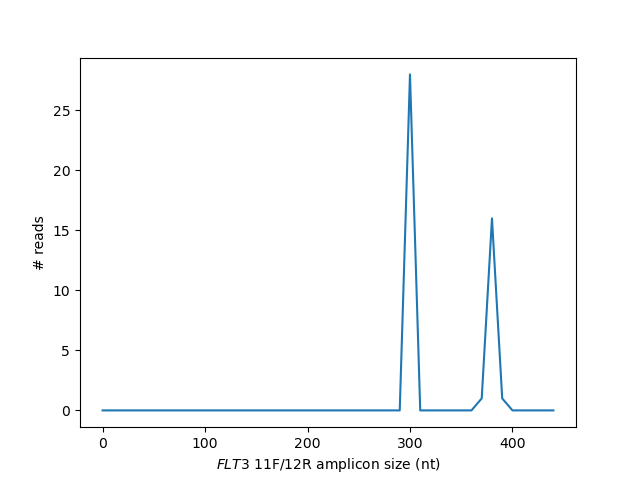

**Figure S2.** Patient samples 0131 and 0133 represent cases of B-ALL with an inconclusive karyotype due to low blast count. (A) For patient sample 0131 (25% blasts), clinical karyotype analysis showed a near haploid karyotype (B) For patient sample 0133 (27% blasts), clinical microarray showed gains of chromosomes 4, 6, 14, 17, 18, and 21.

(A)

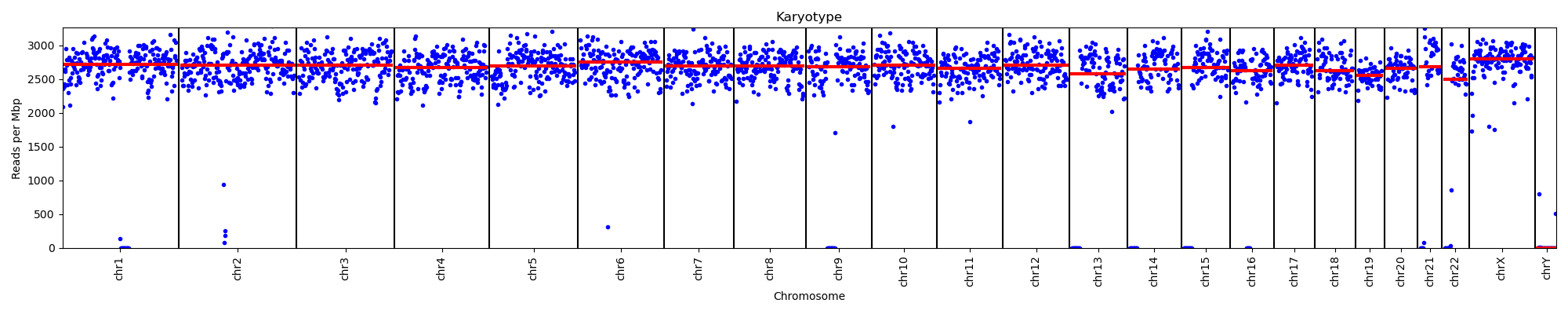

(B)
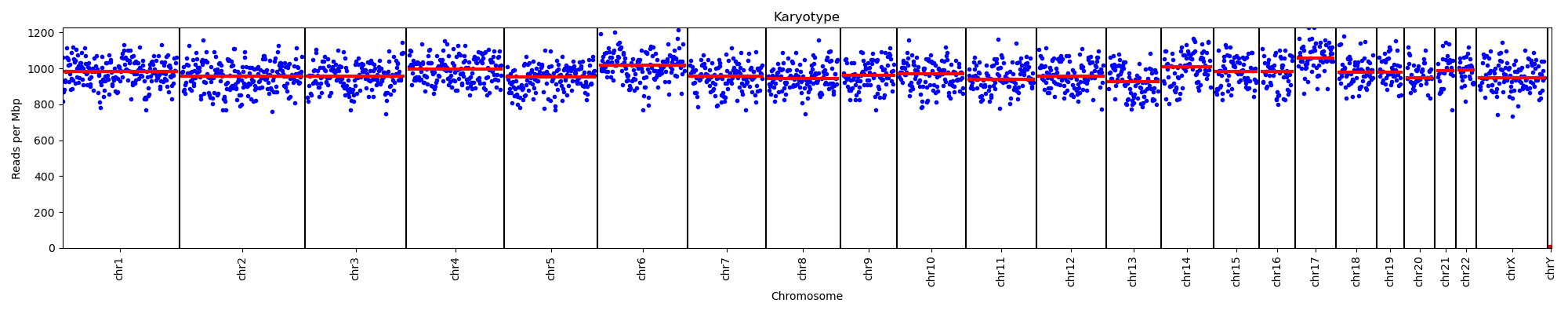

**Figure S3.** Sample 0164 with iAMP21. Depth of coverage over chromosome 21 (15Mbp - 45Mbp) shows segmental copy number variation with six apparent copies over the region spanning *RUNX1*.

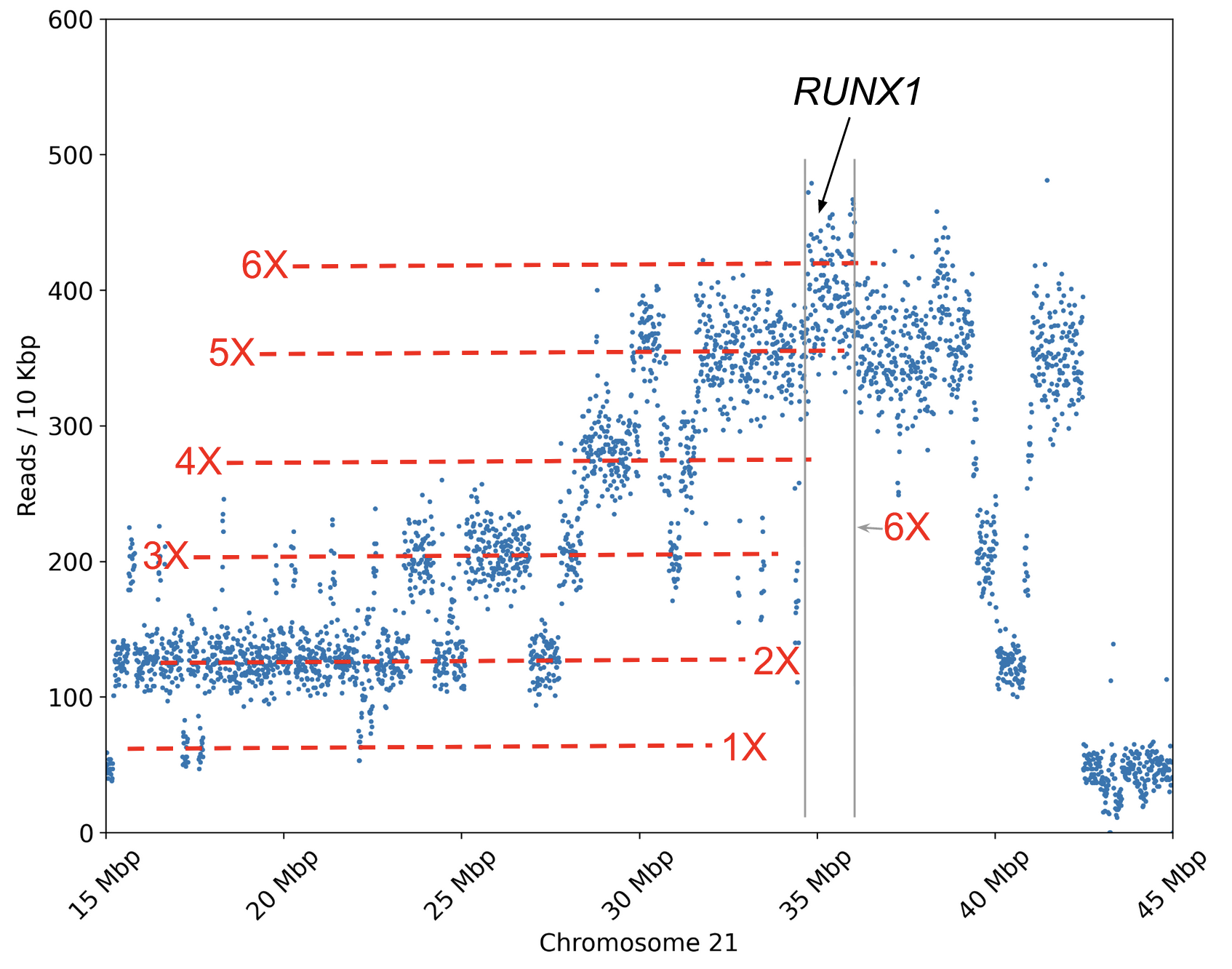

**Figure S4**. Sequencing depth profiles illustrating karyotype variation at early and late timepoints. Sample 0227 depth profile is shown when it is called low hypodiploid after 2 minutes (A) and at the end of sequencing (B). Sample 0173 is shown when it is called high hyperdiploid after 5 minutes (C) and at the end of sequencing (D).

(A)

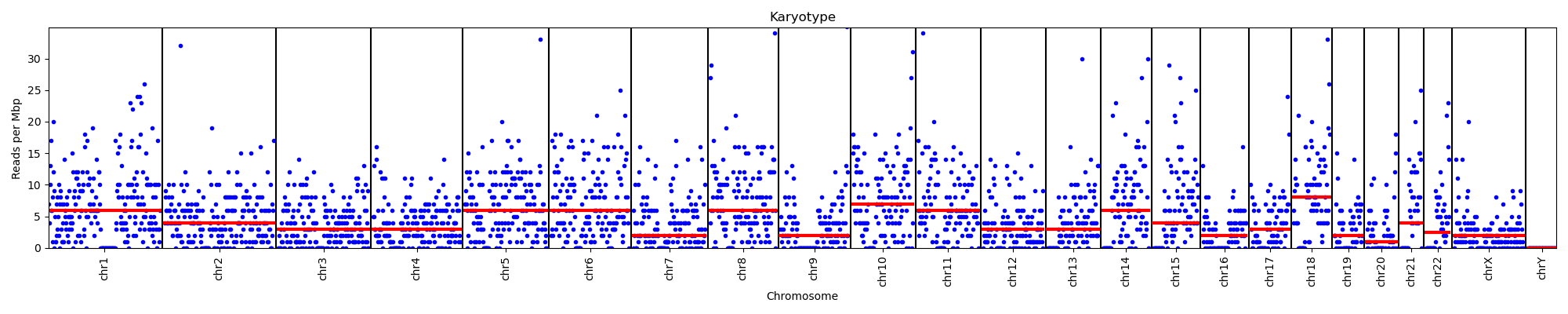

(B)

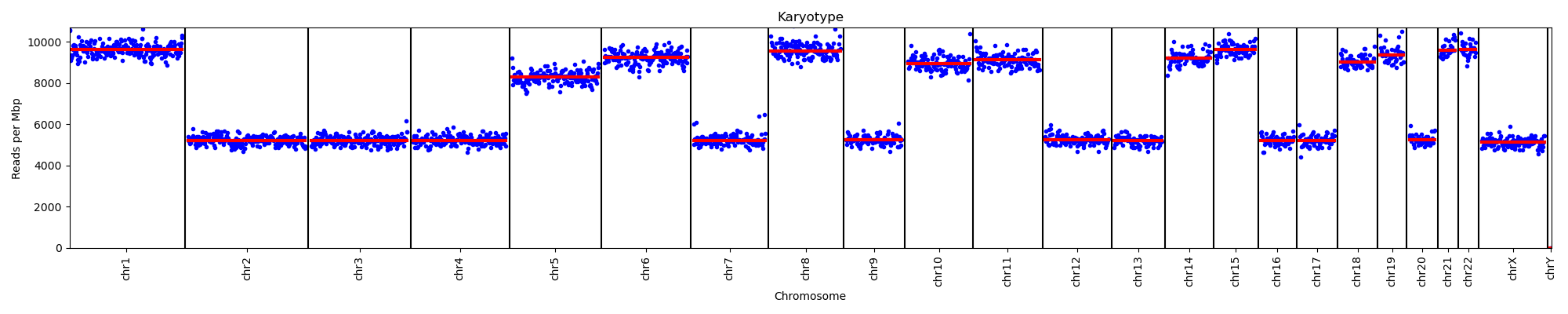

(C)

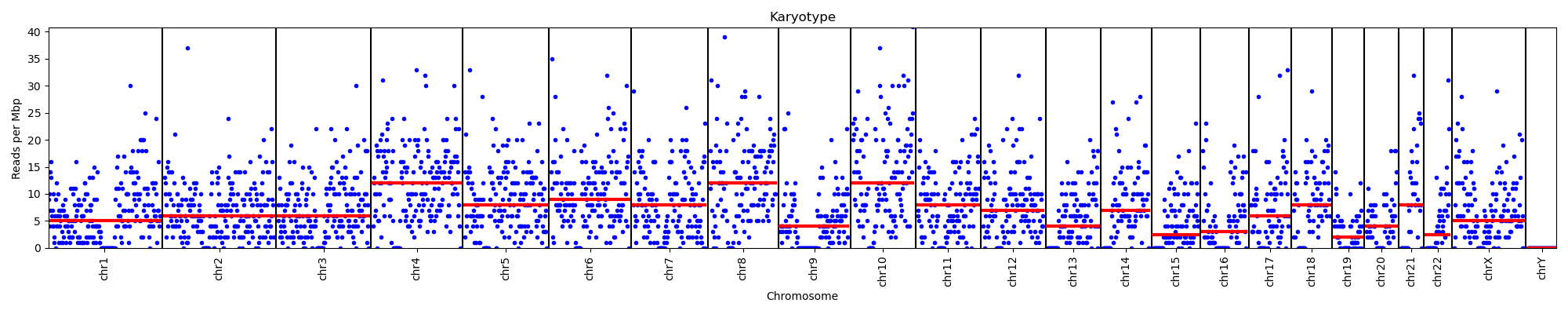

(D)

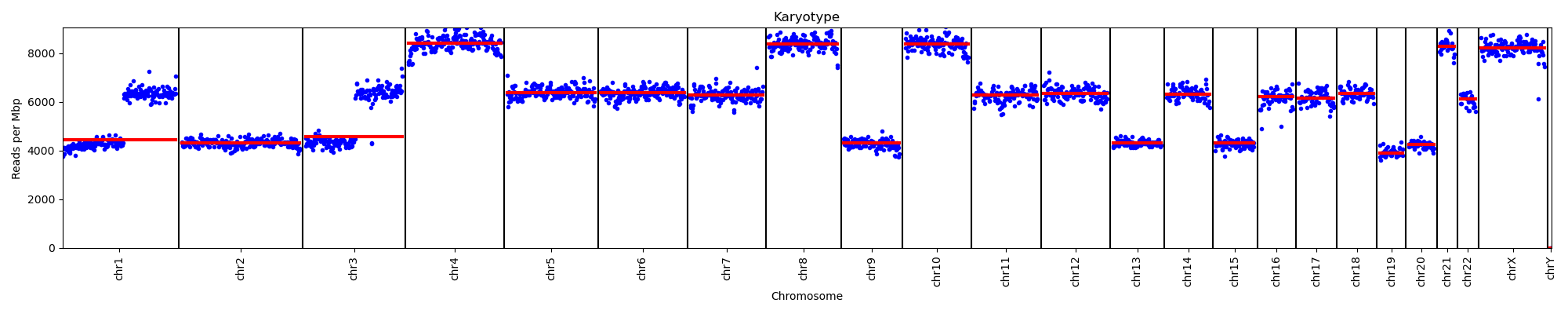

**Figure S5.** Genome-wide sequencing depth profiles for all samples showing karyotype variation

Nano_Leuk_0050

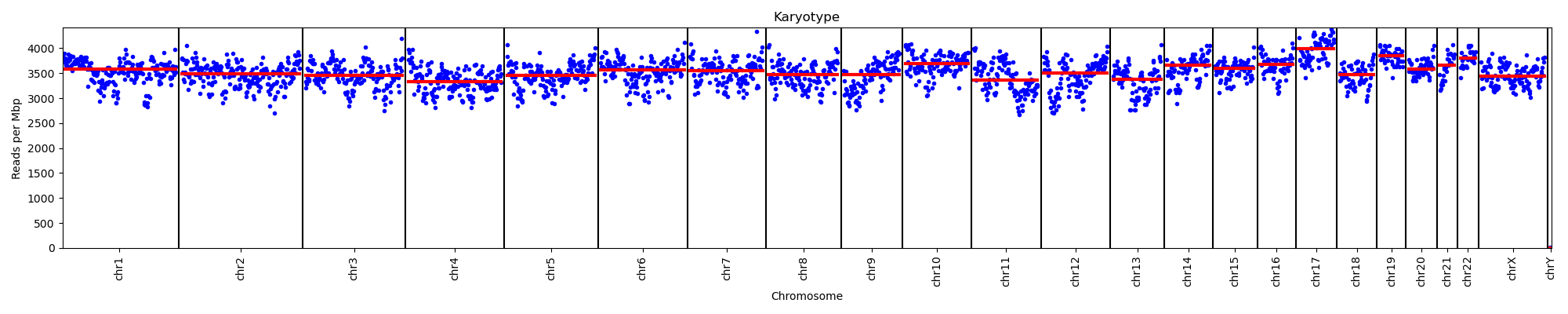

Nano_Leuk_0052
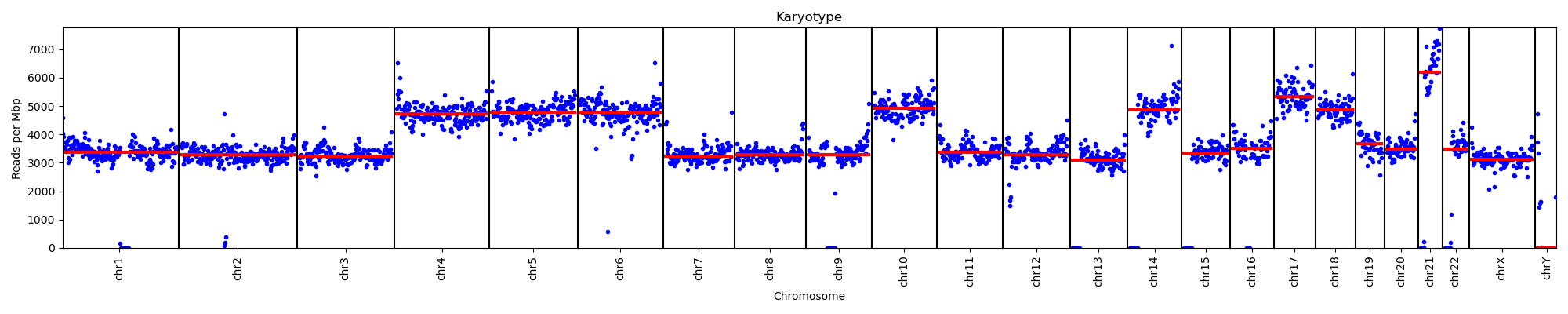

Nano_Leuk_0055

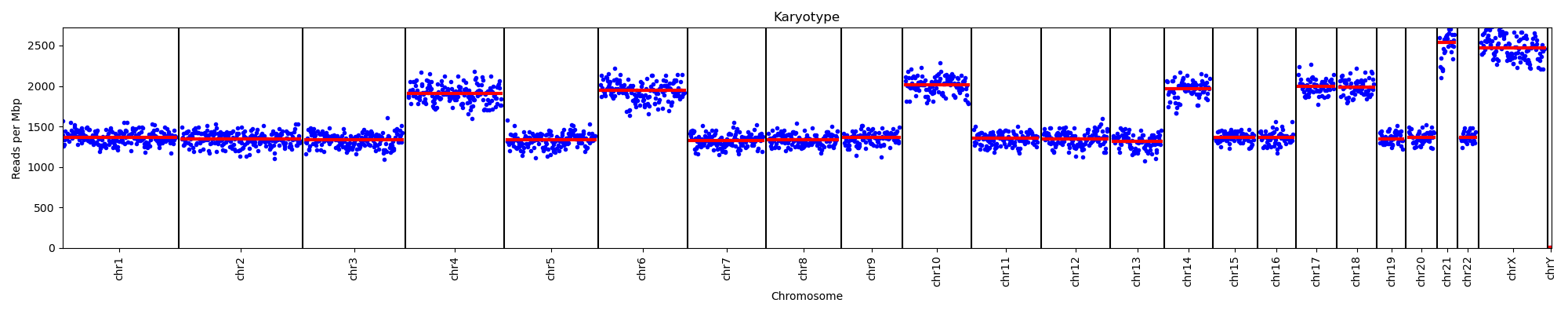

Nano_Leuk_0057

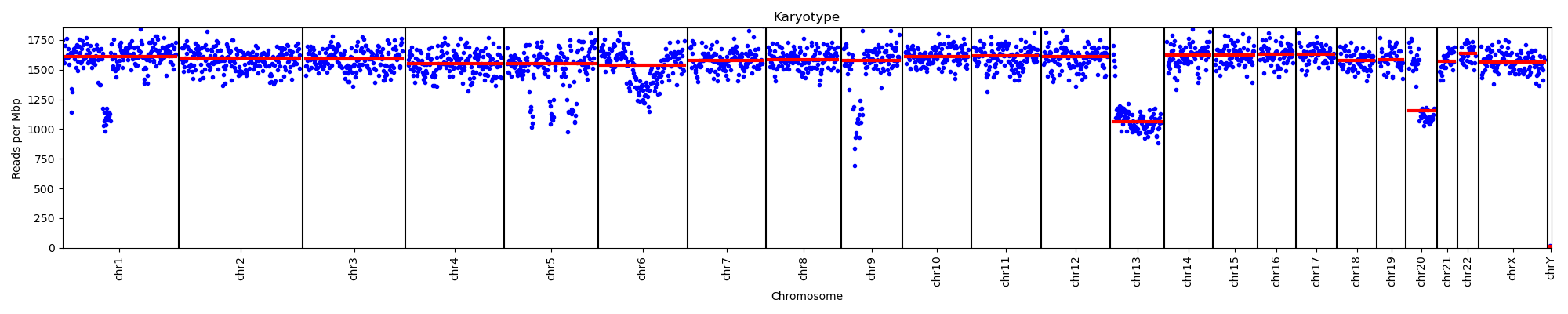

Nano_Leuk_0058

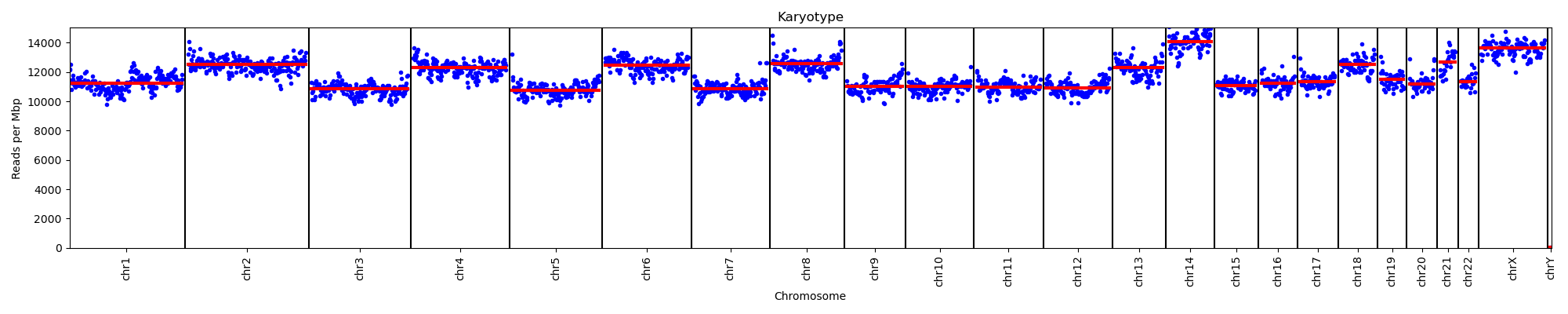

Nano_Leuk_0060

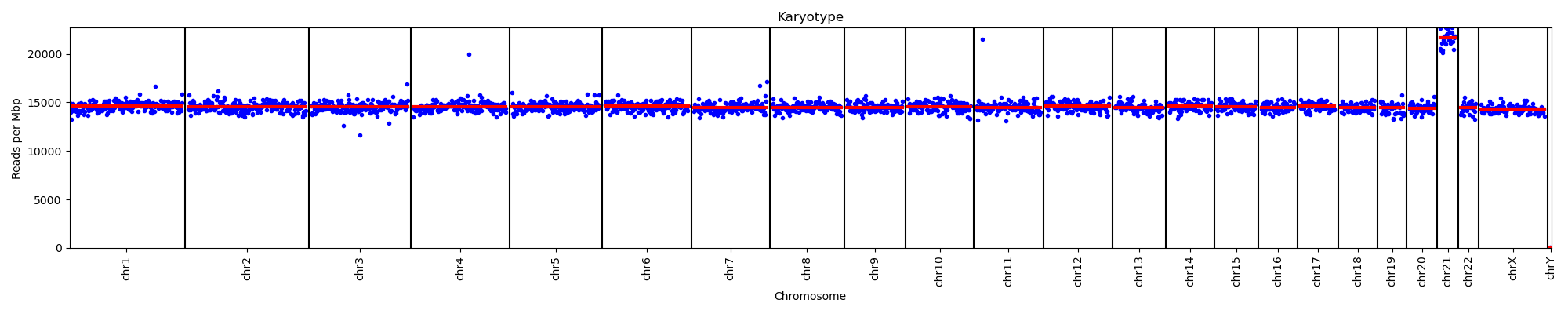

Nano_Leuk_0129
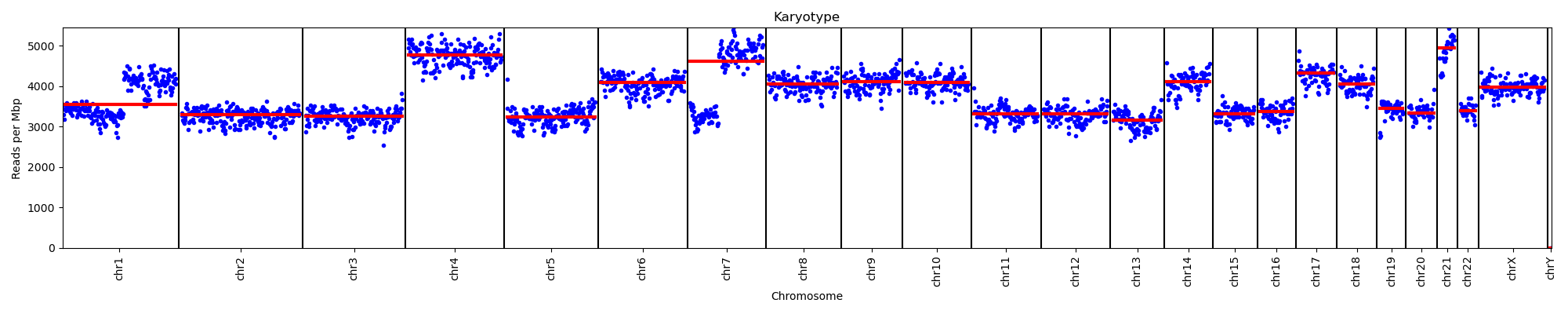

Nano_Leuk_0130

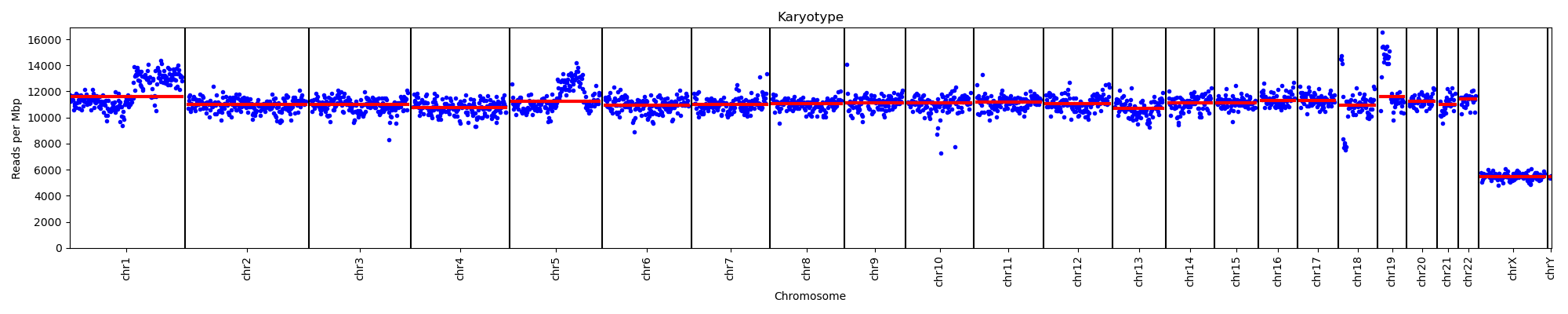

Nano_Leuk_0131
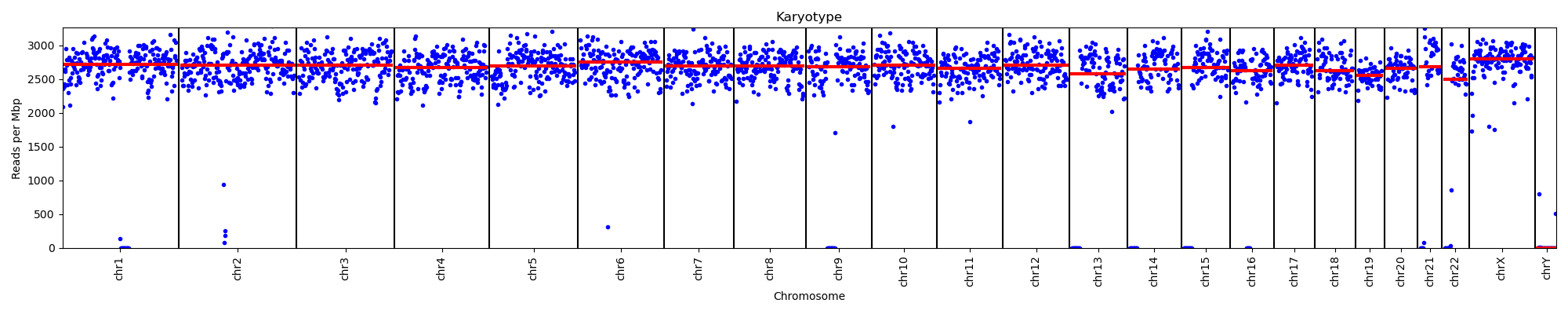

Nano_Leuk_0132
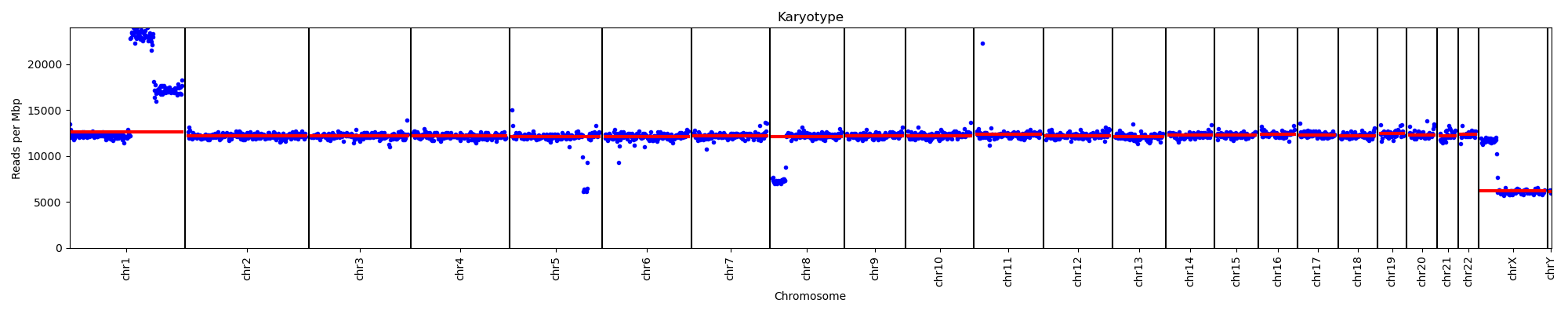

Nano_Leuk_0133
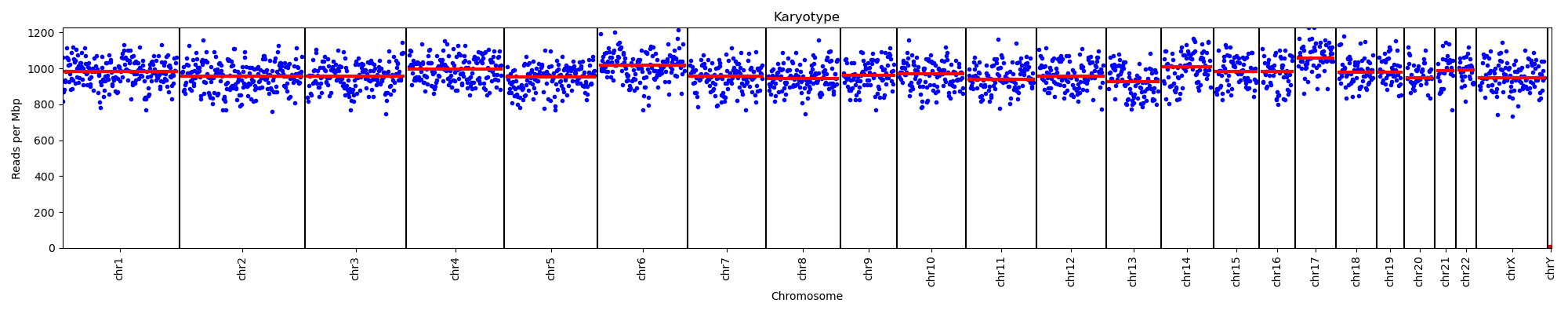

Nano_Leuk_0136

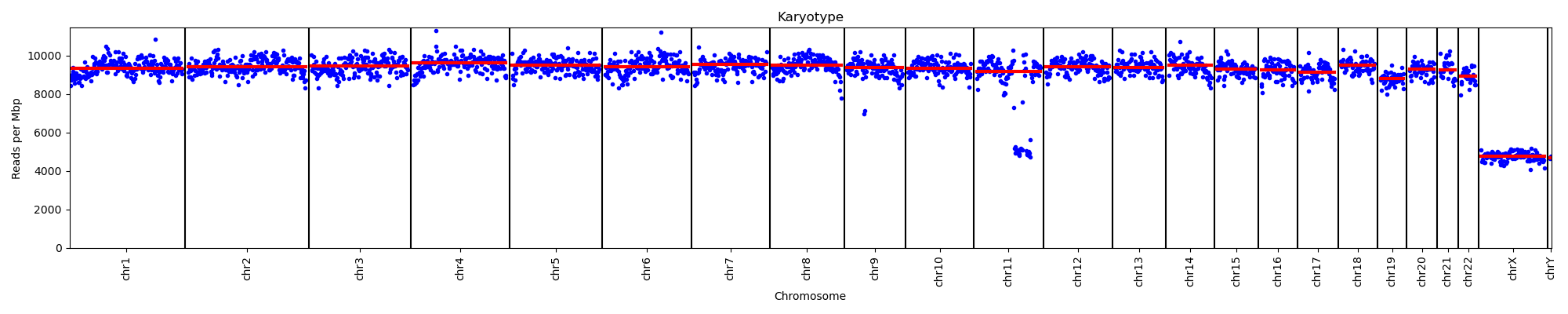

Nano_Leuk_0140

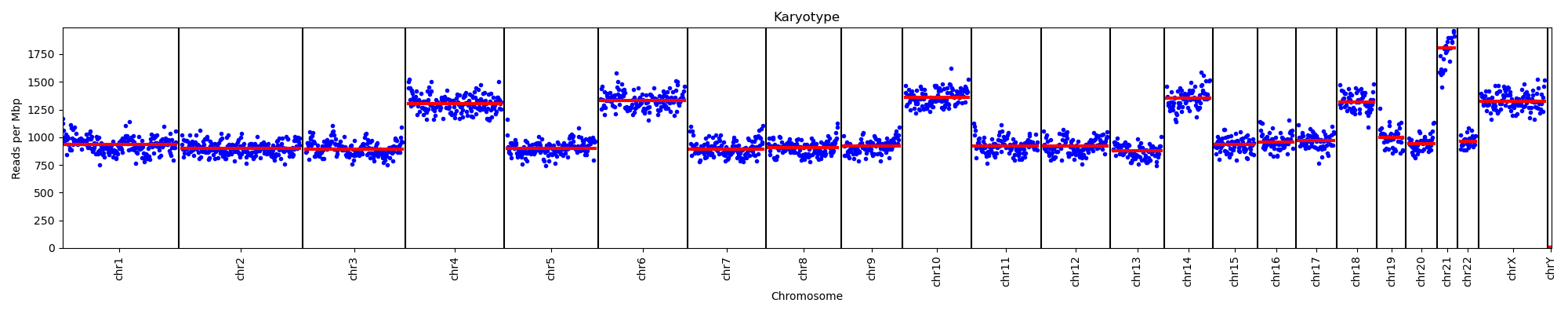

Nano_Leuk_0141

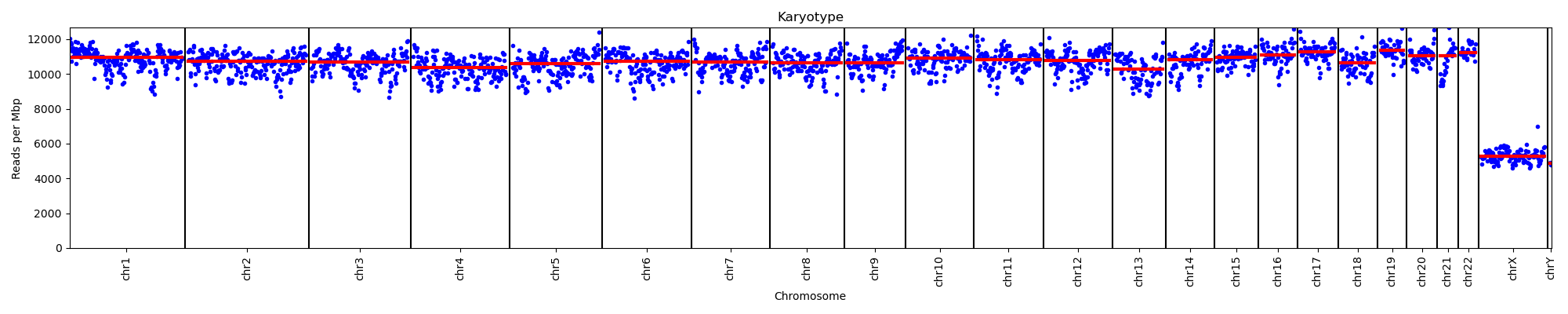

Nano_Leuk_0145
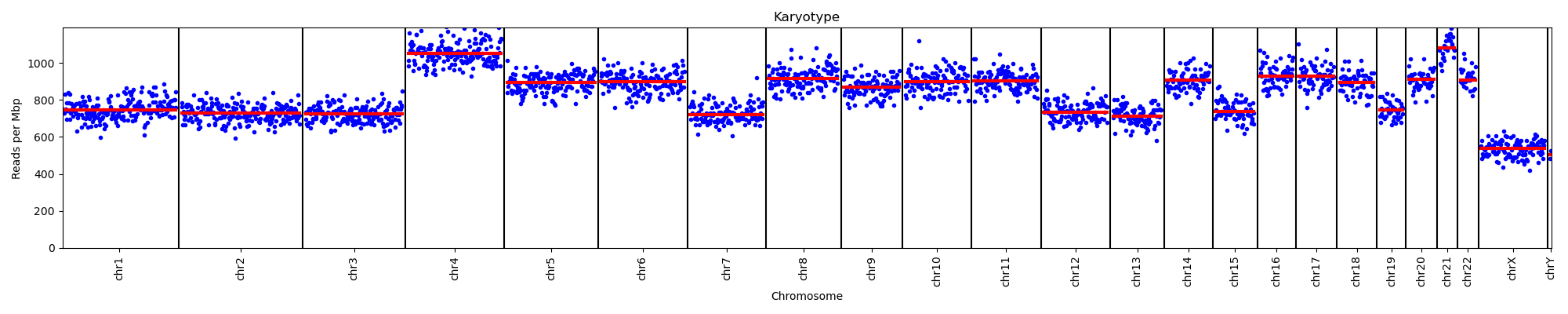

Nano_Leuk_0148
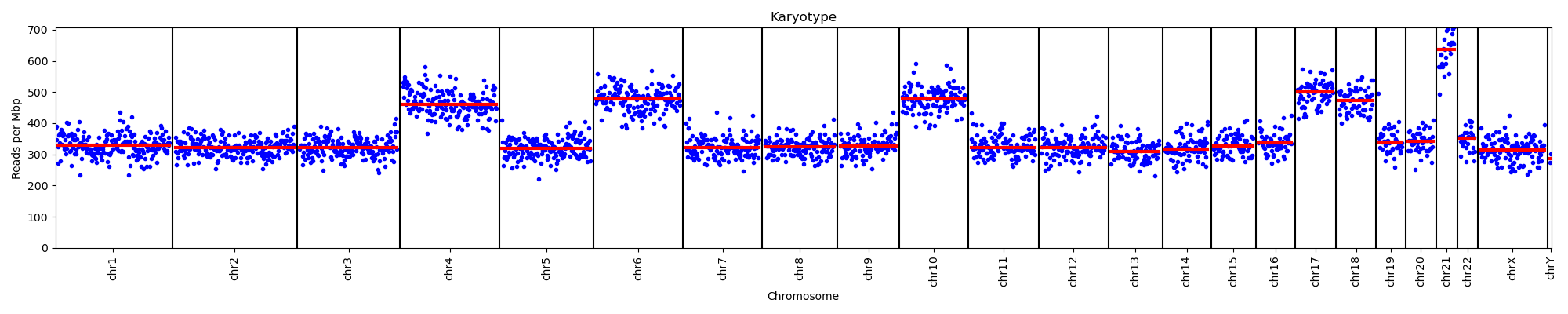

Nano_Leuk_0149

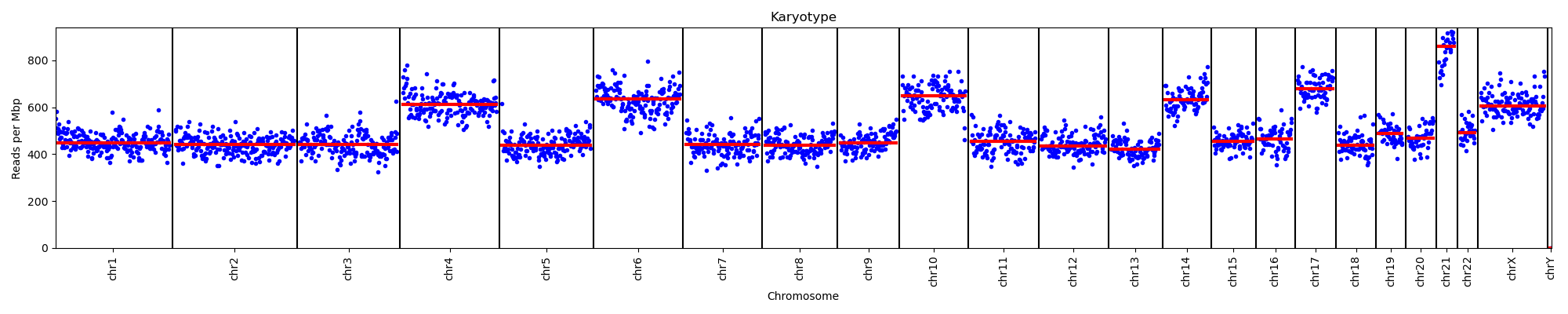

Nano_Leuk_0151
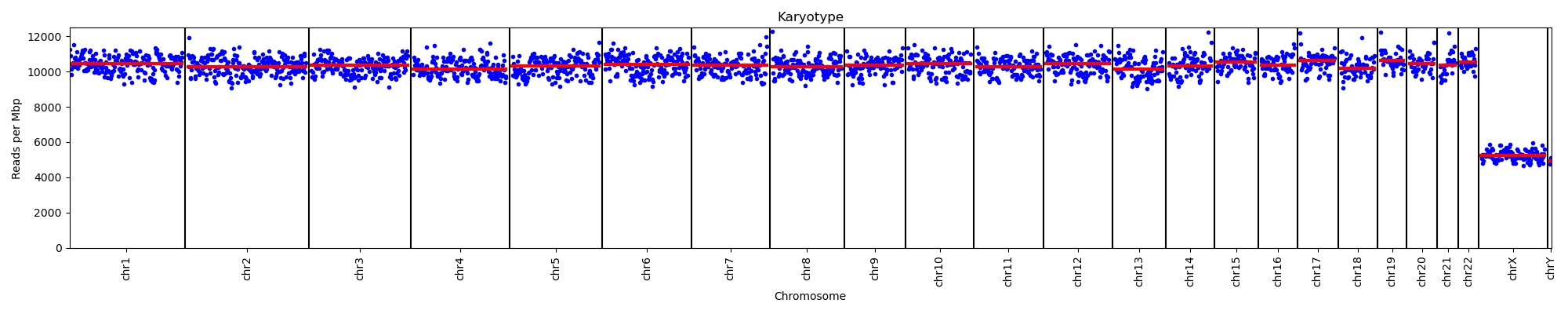

Nano_Leuk_0152
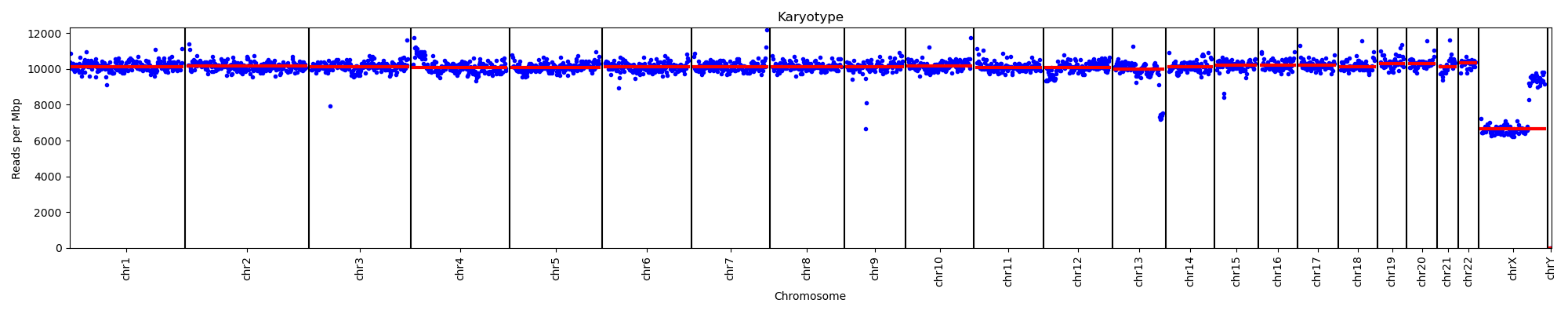

Nano_Leuk_0153
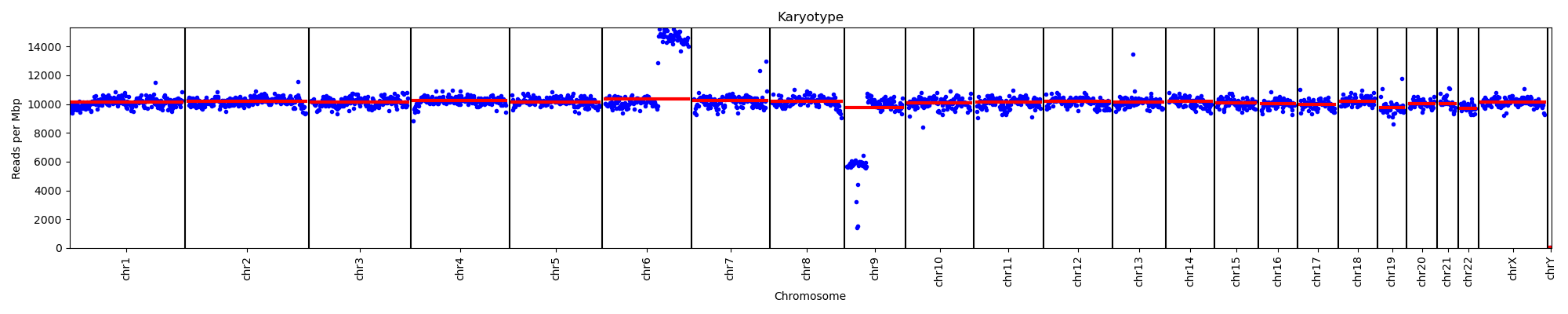

Nano_Leuk_0154
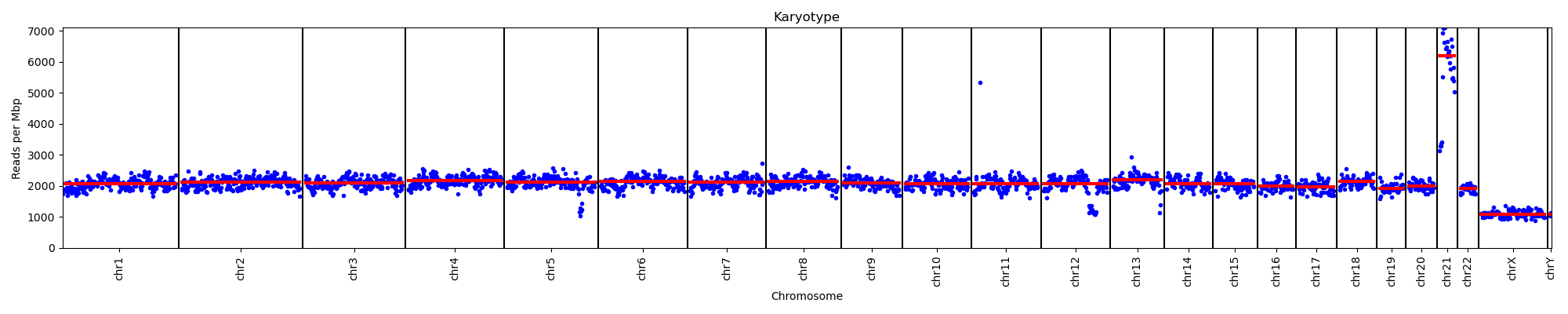

Nano_Leuk_0155
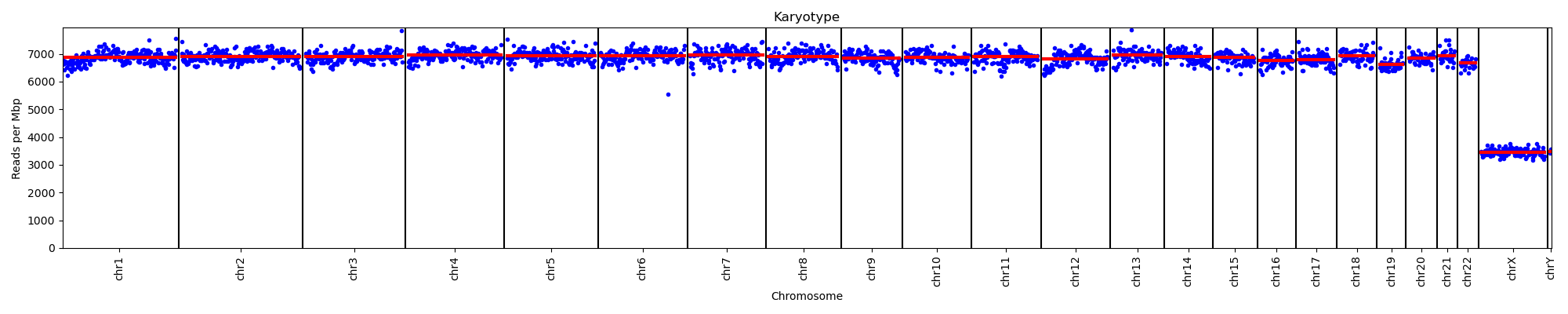

Nano_Leuk_0156
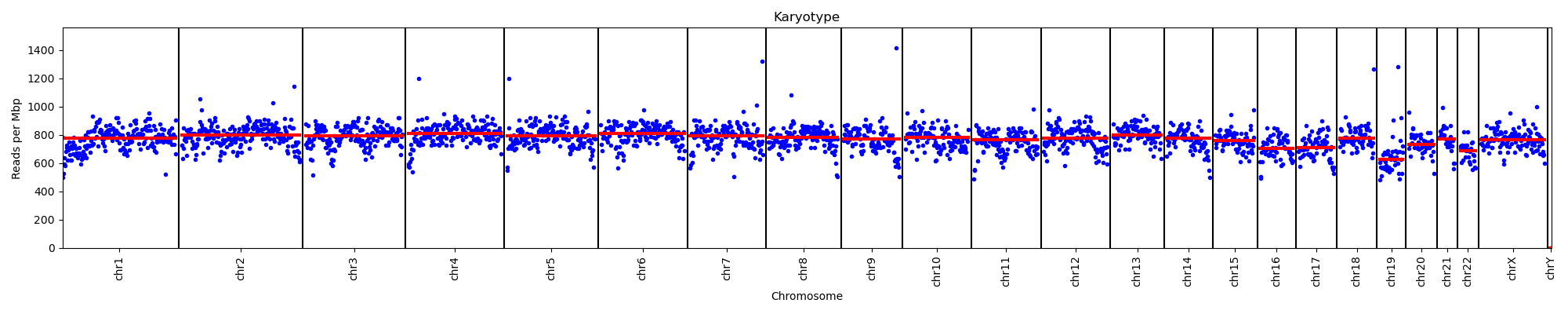

Nano_Leuk_0157

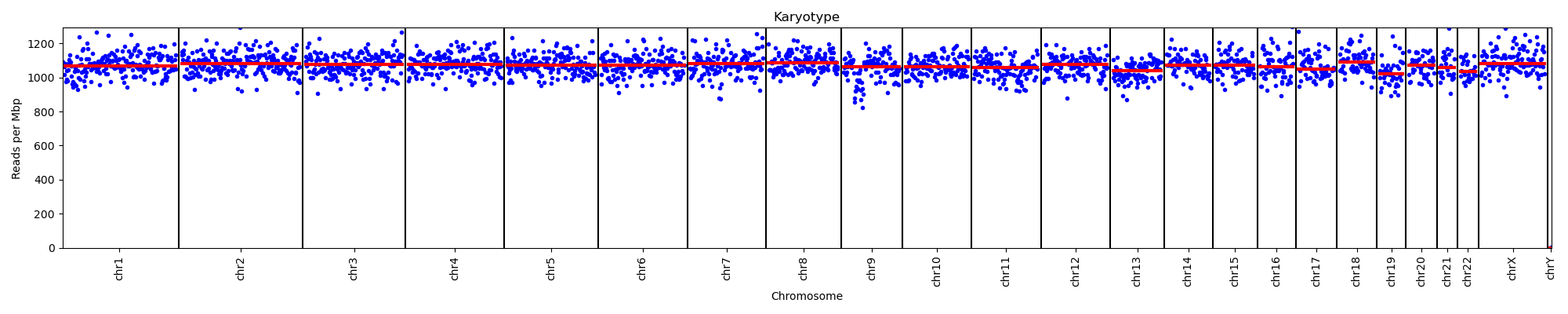

Nano_Leuk_0158

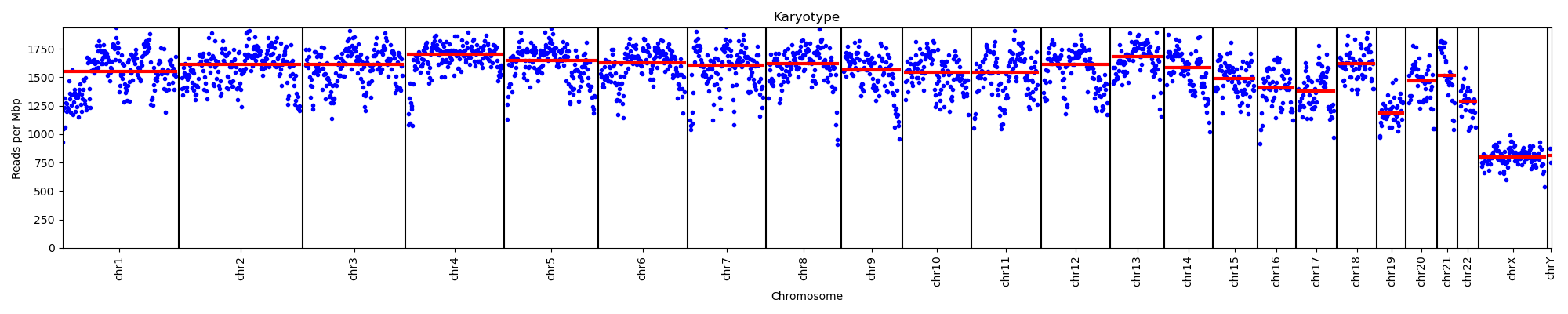

Nano_Leuk_0159
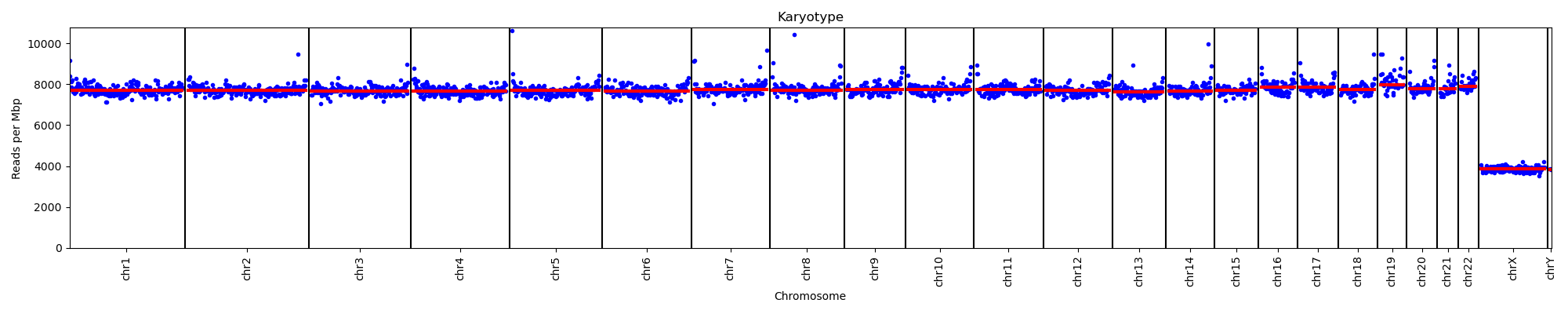

Nano_Leuk_0161
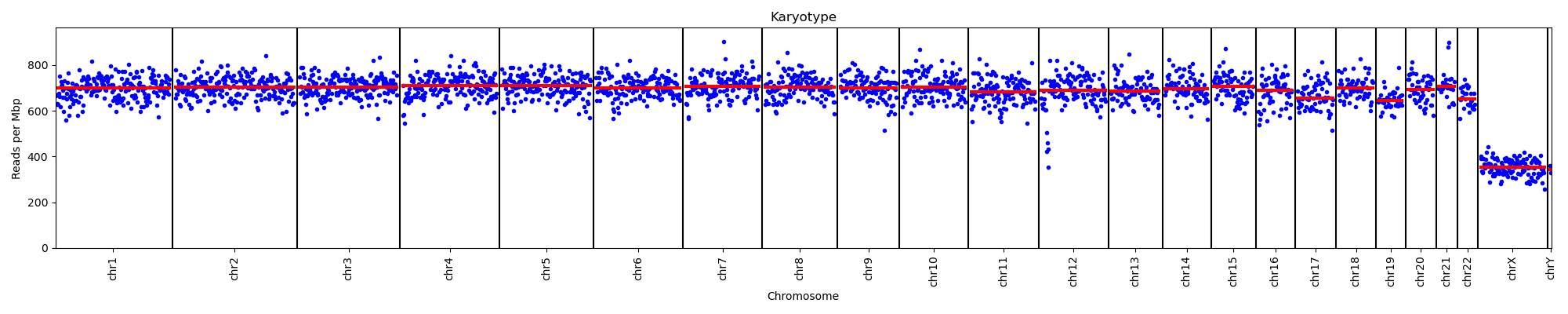

Nano_Leuk_0162
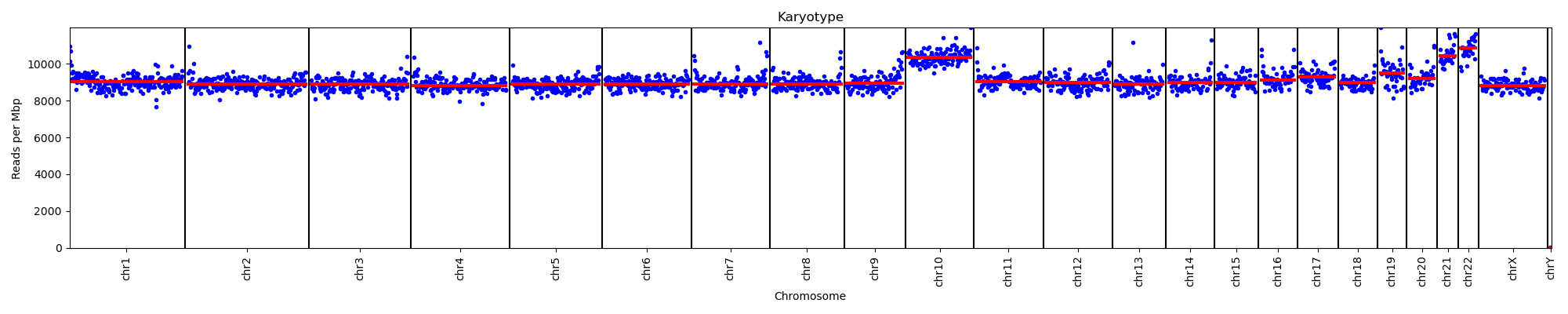

Nano_Leuk_0163

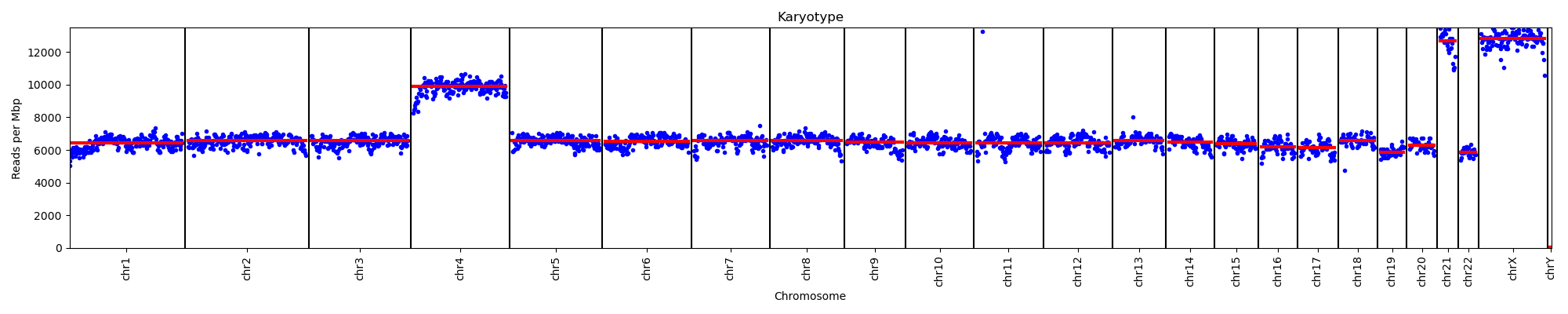

Nano_Leuk_0164
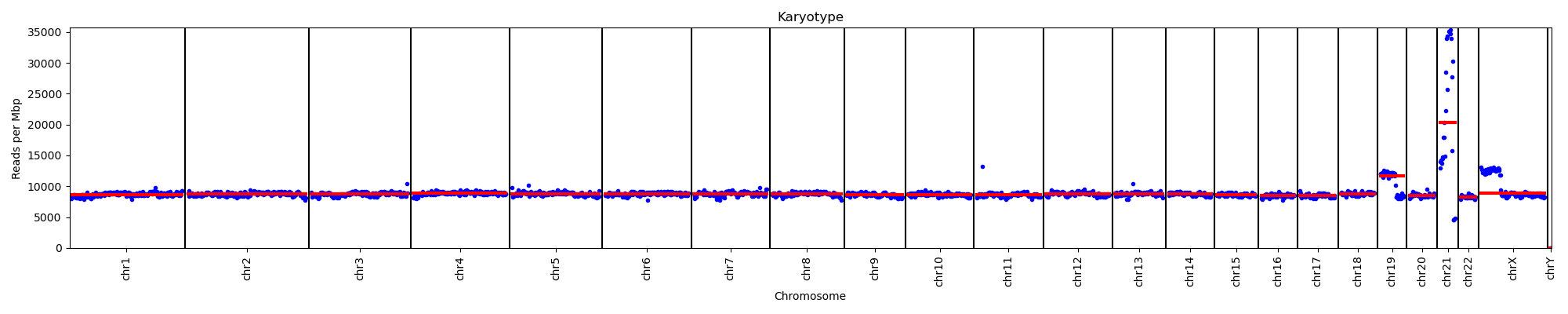

Nano_Leuk_0166
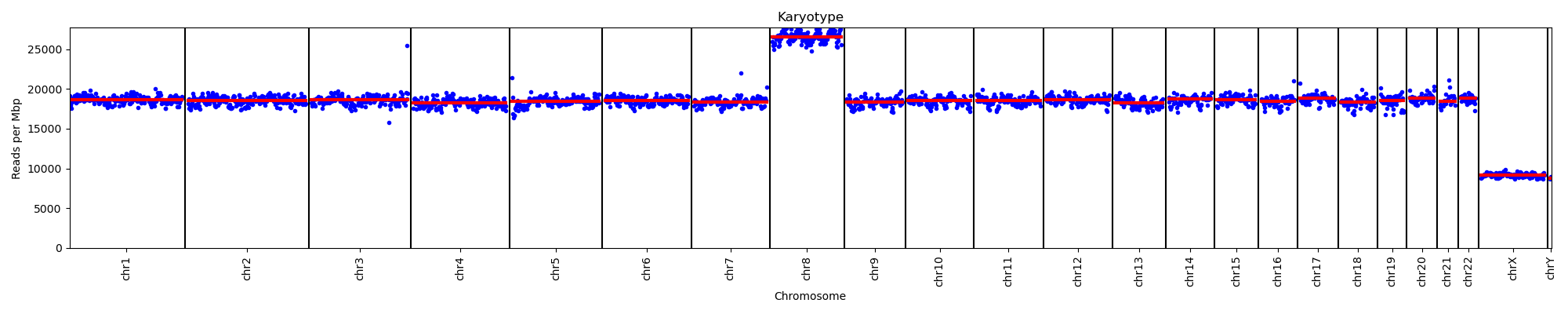

Nano_Leuk_0168

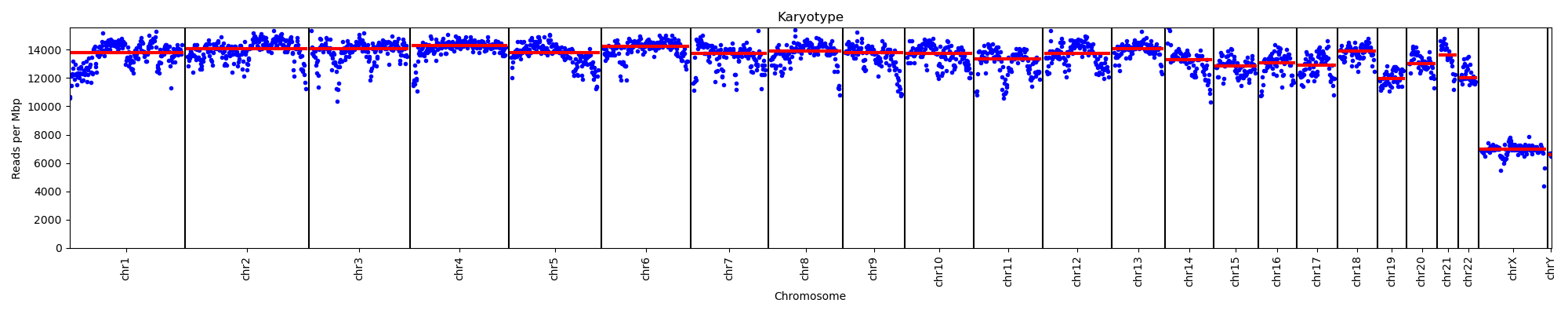

Nano_Leuk_0170

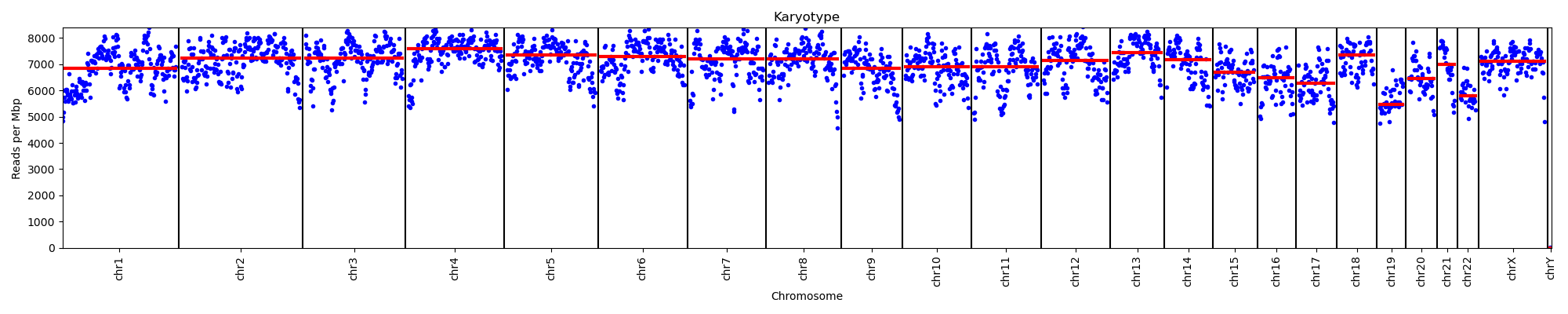

Nano_Leuk_0171
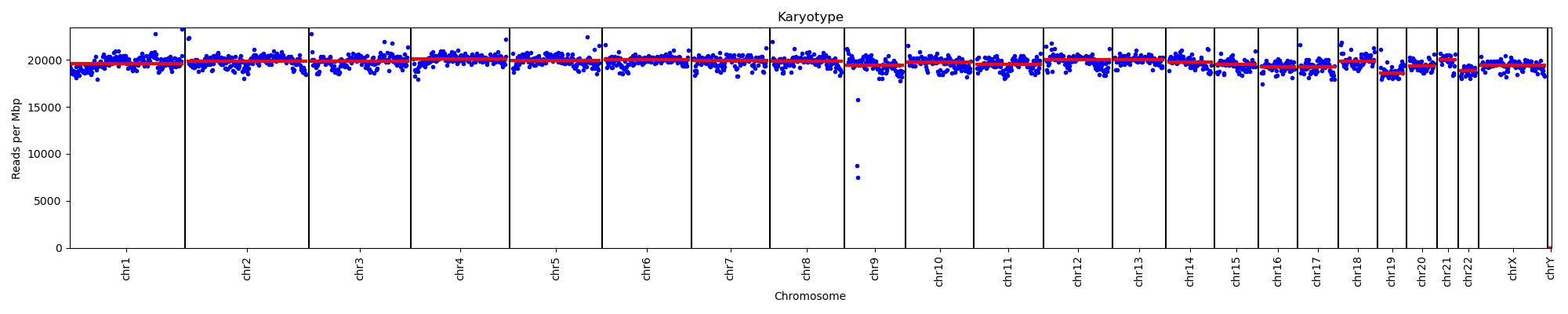

Nano_Leuk_0172
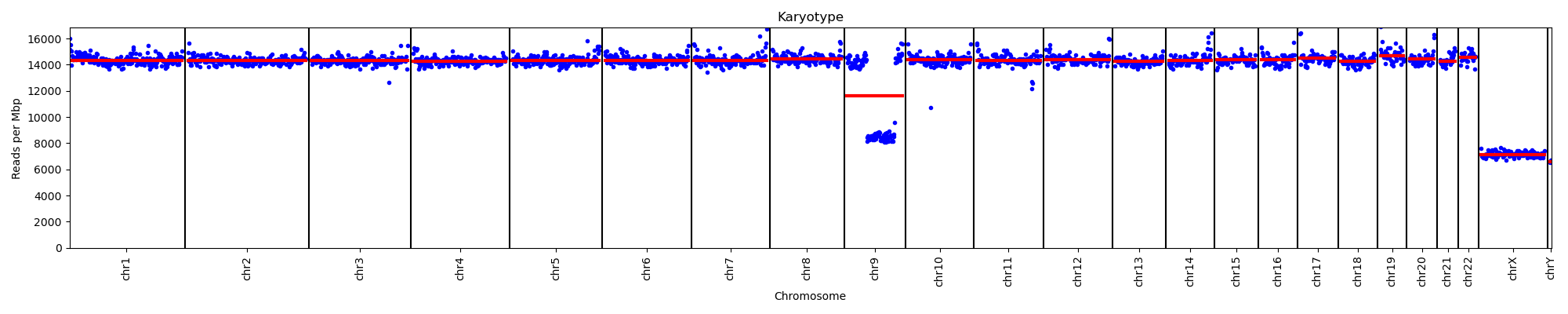

Nano_Leuk_0173
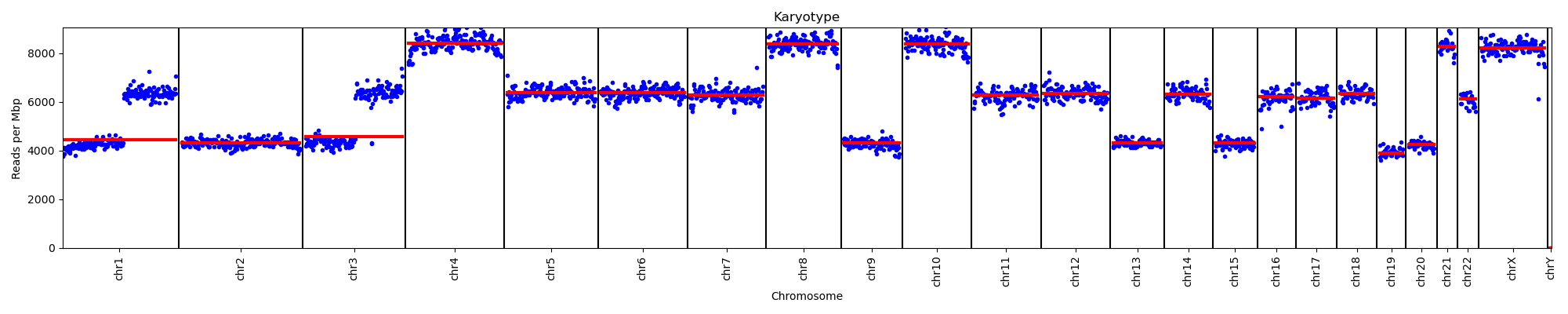

Nano_Leuk_0203
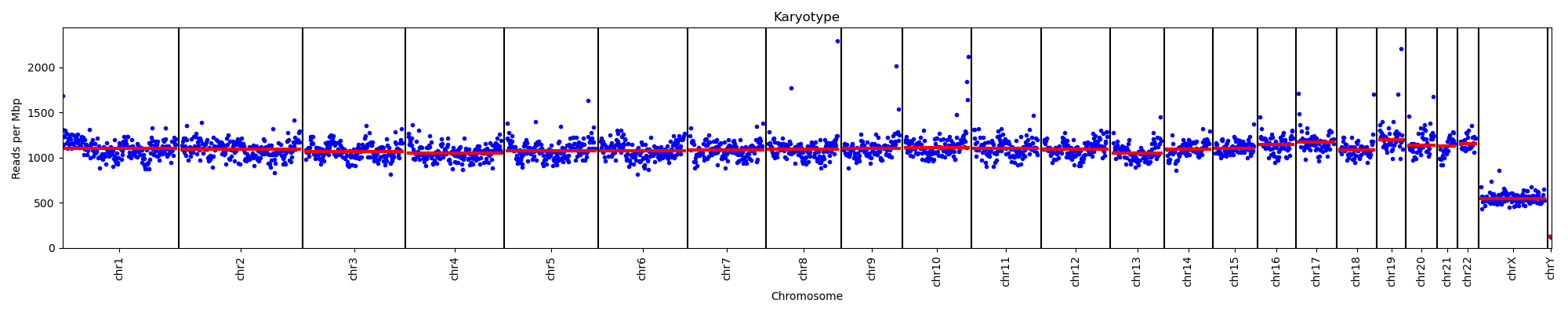

Nano_Leuk_0209
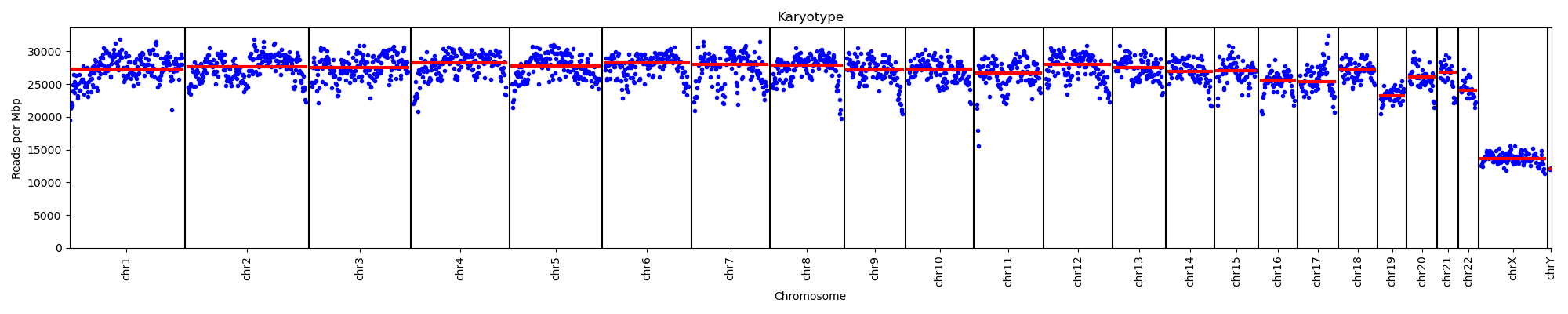

Nano_Leuk_0211
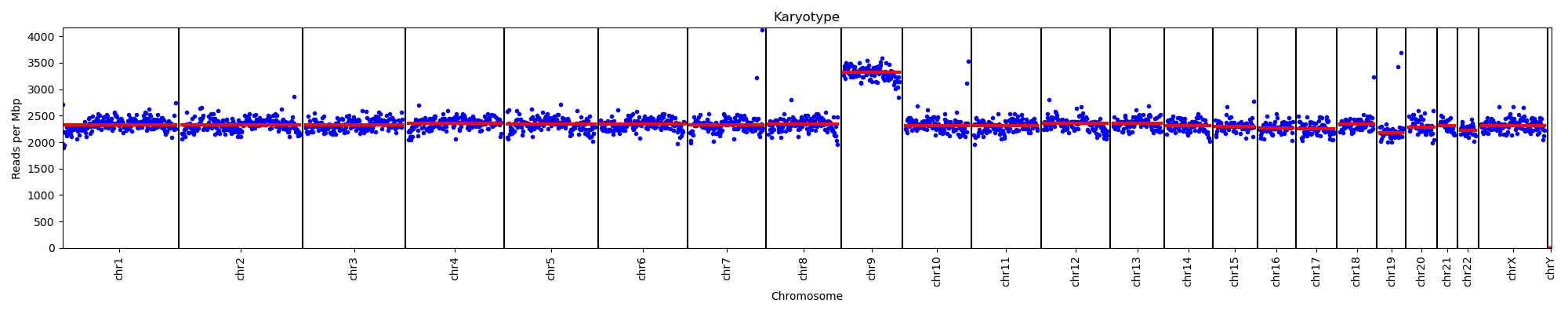

Nano_Leuk_0222

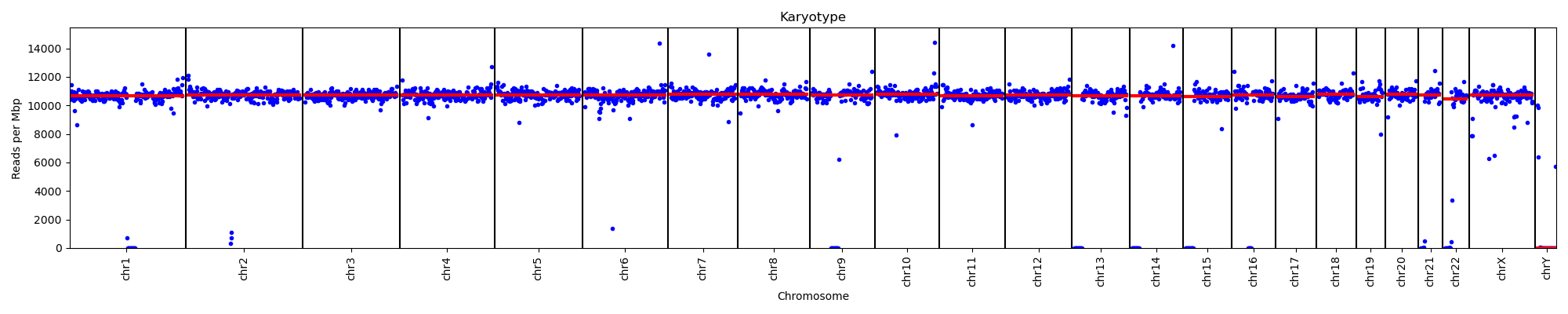

Nano_Leuk_0223
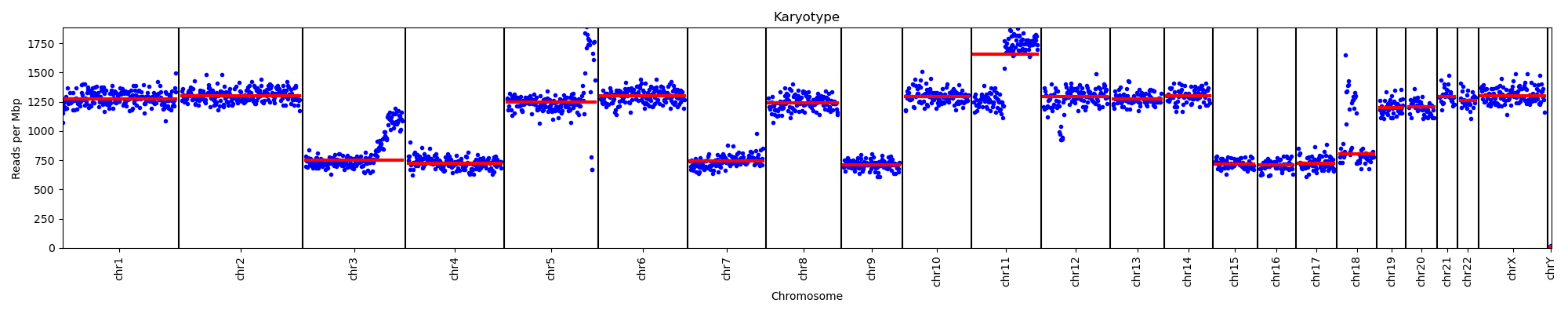

Nano_Leuk_0224
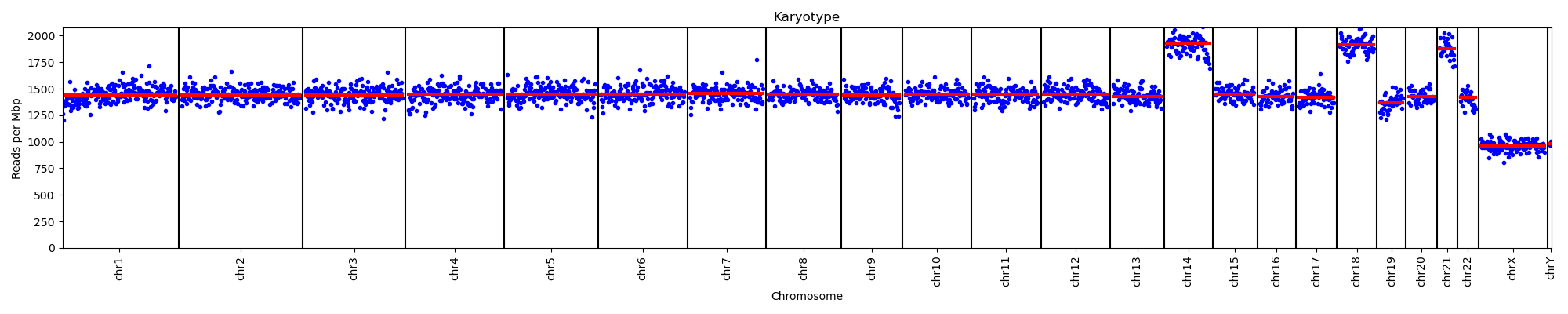

Nano_Leuk_0225

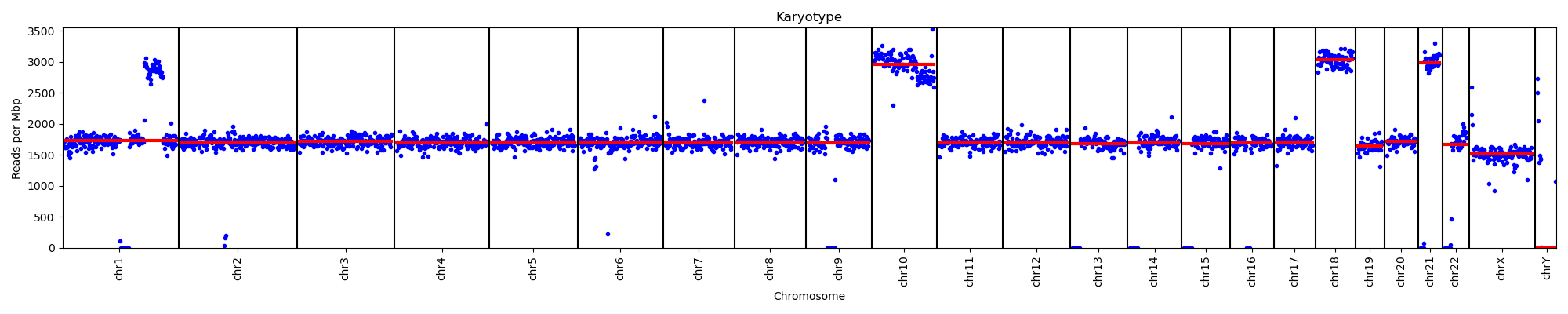

Nano_Leuk_0226
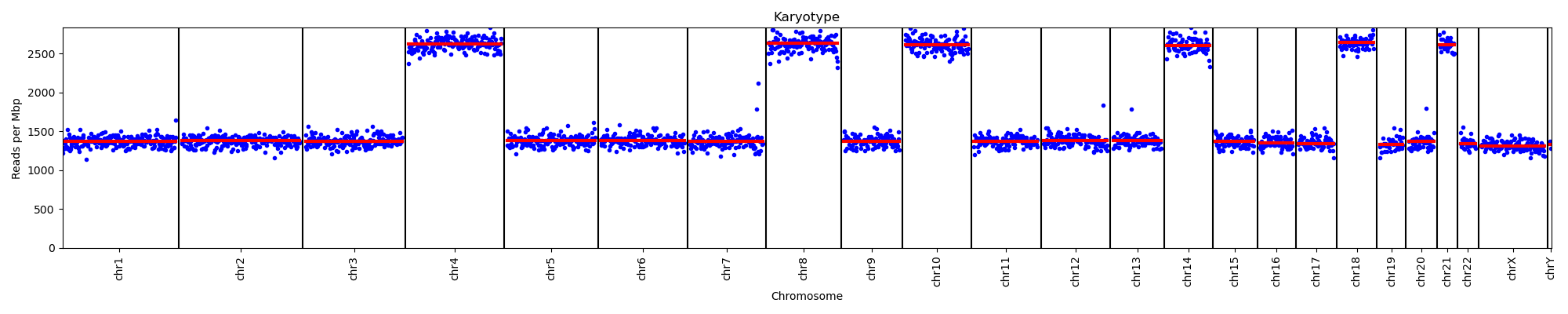

Nano_Leuk_0227
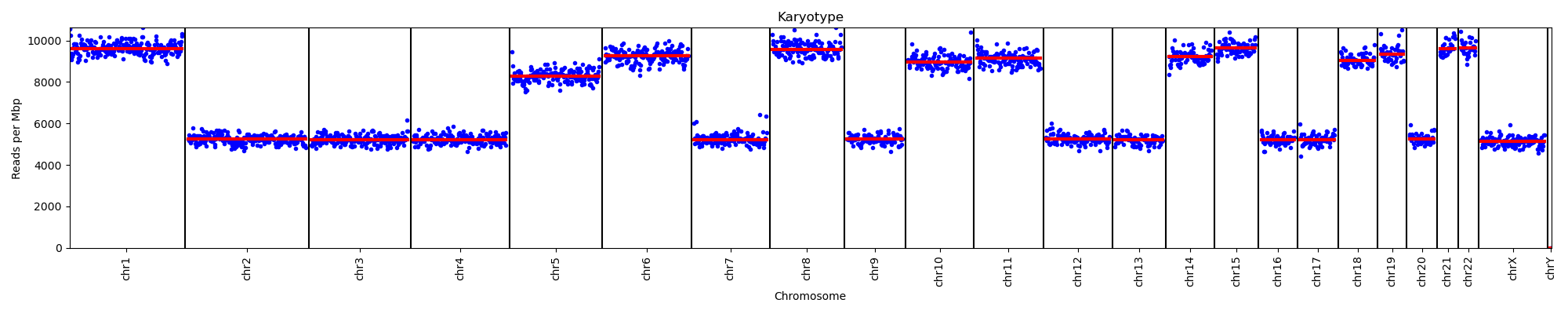

Nano_Leuk_0228
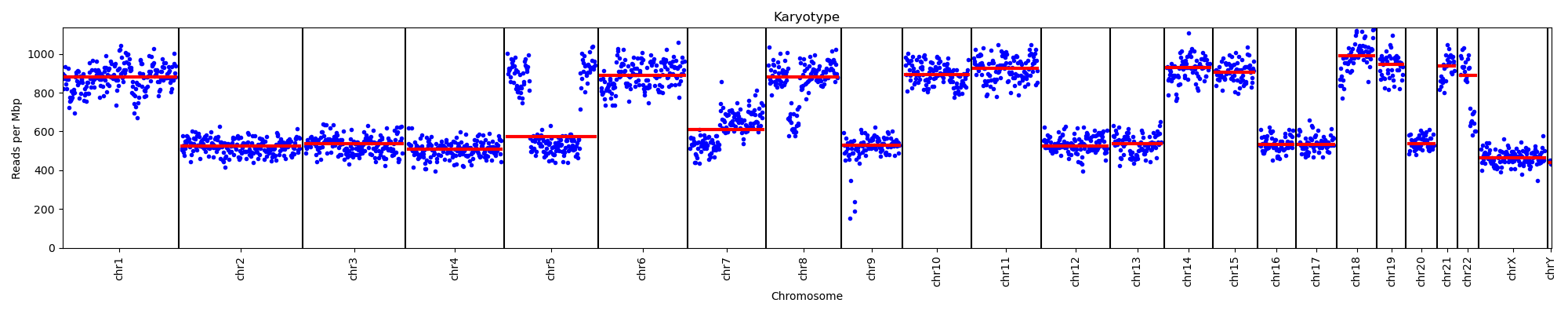

Nano_Leuk_0229
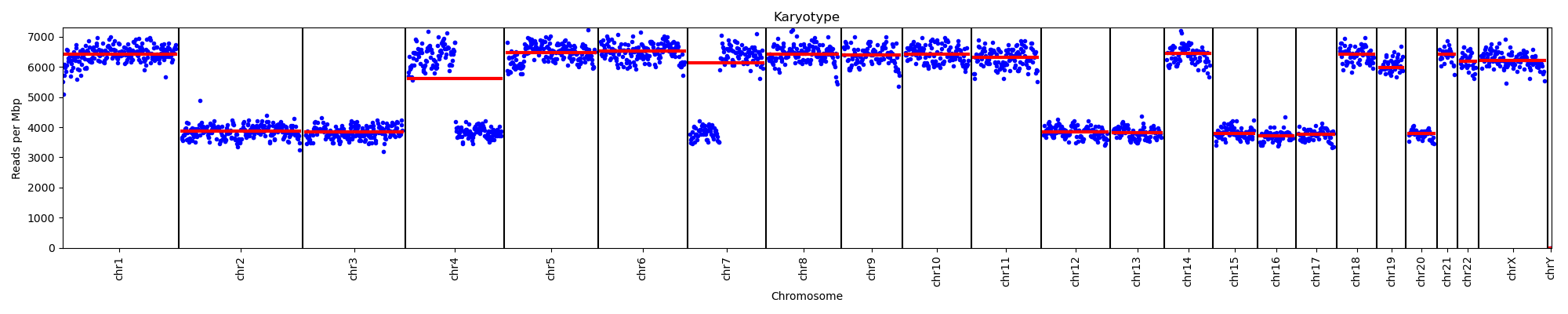

Nano_Leuk_0230
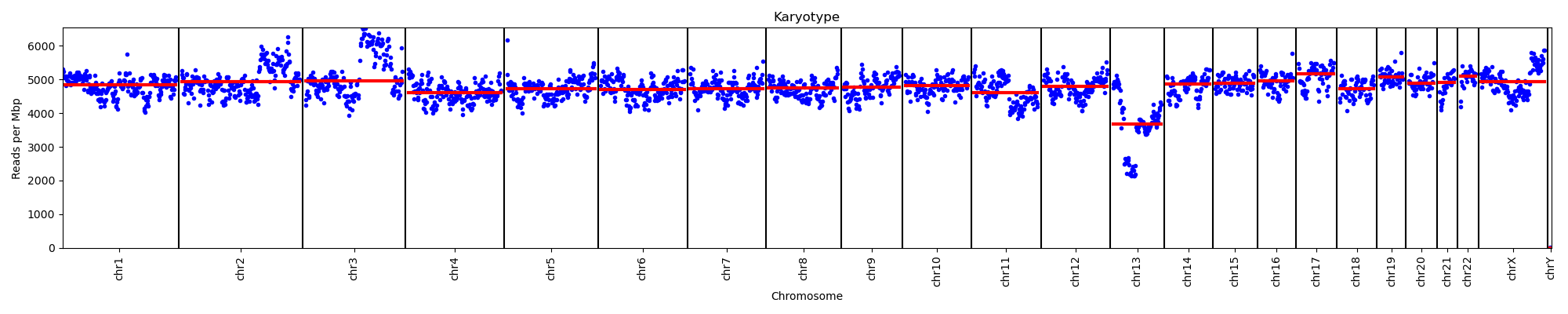

Nano_Leuk_0231
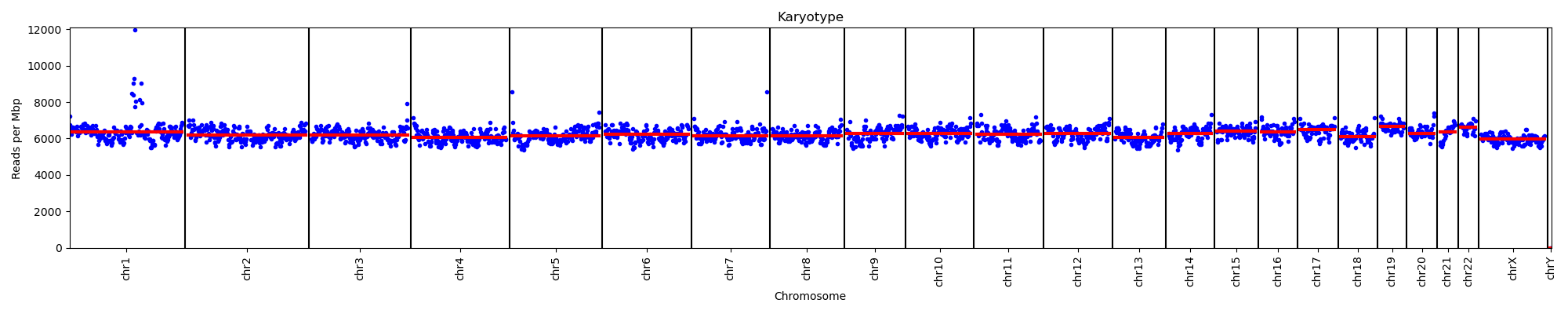

Nano_Leuk_0233
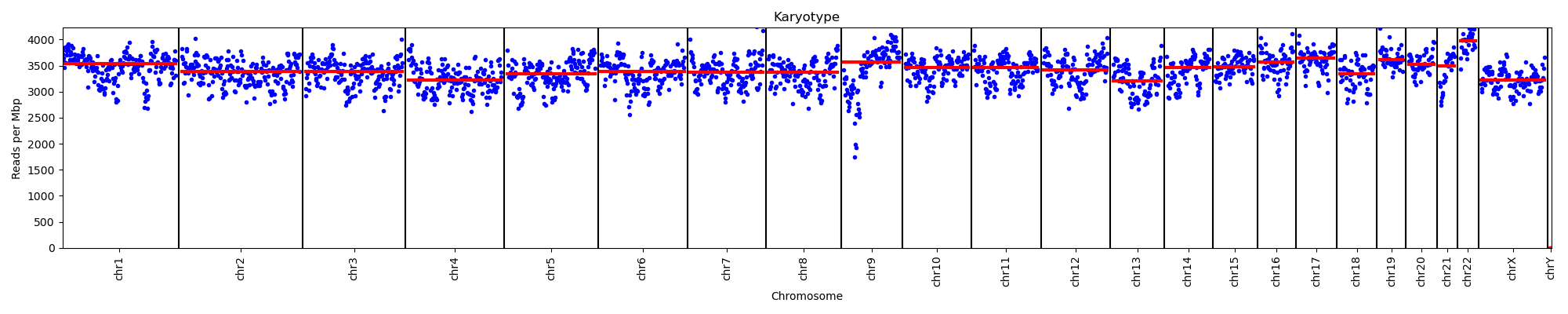

Nano_Leuk_0234
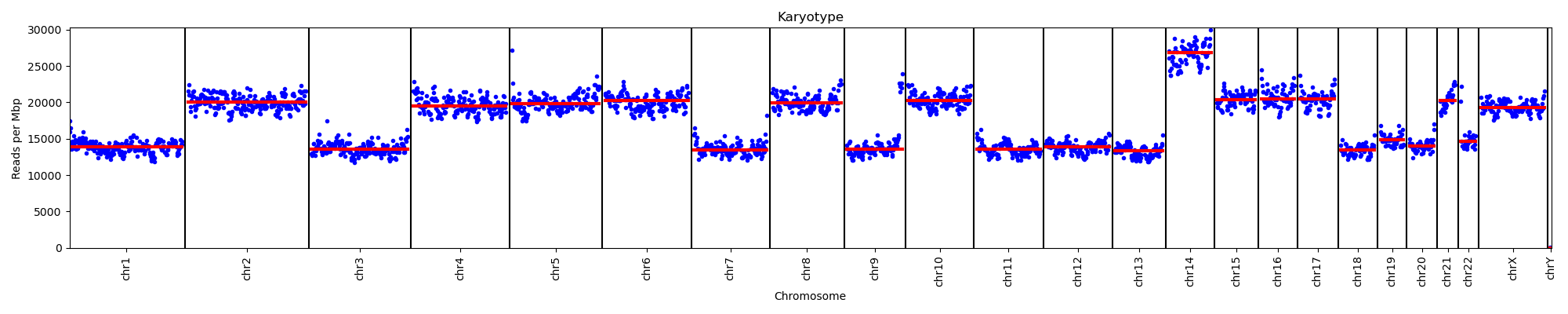

Nano_Leuk_0238
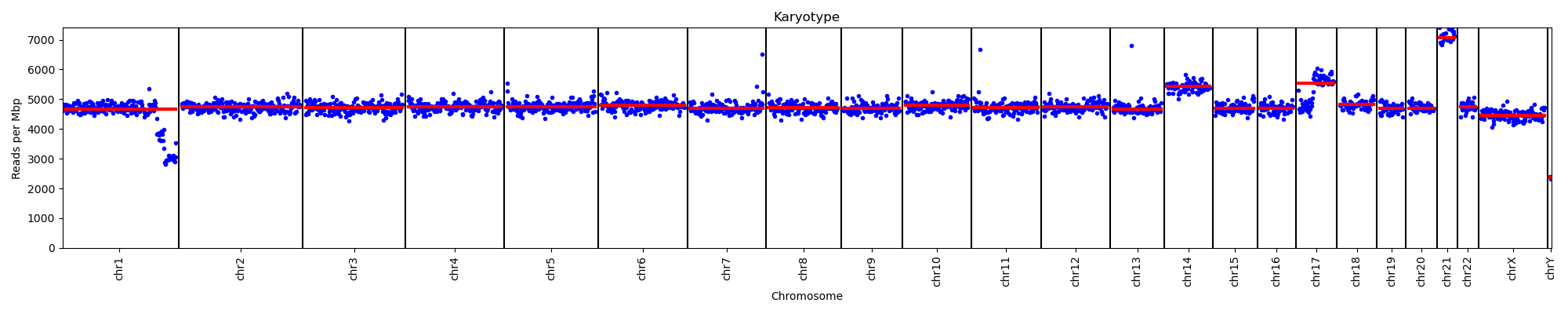

Nano_Leuk_0240
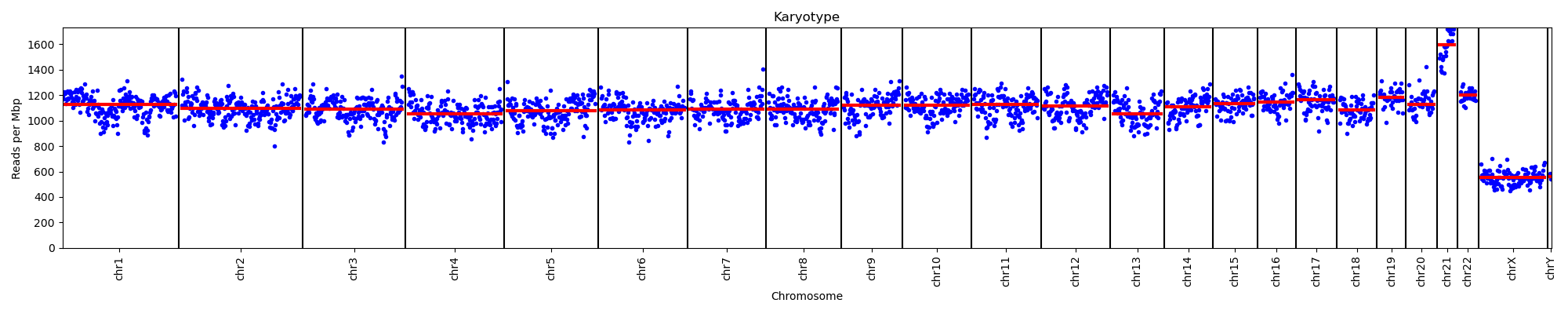

Nano_Leuk_0242
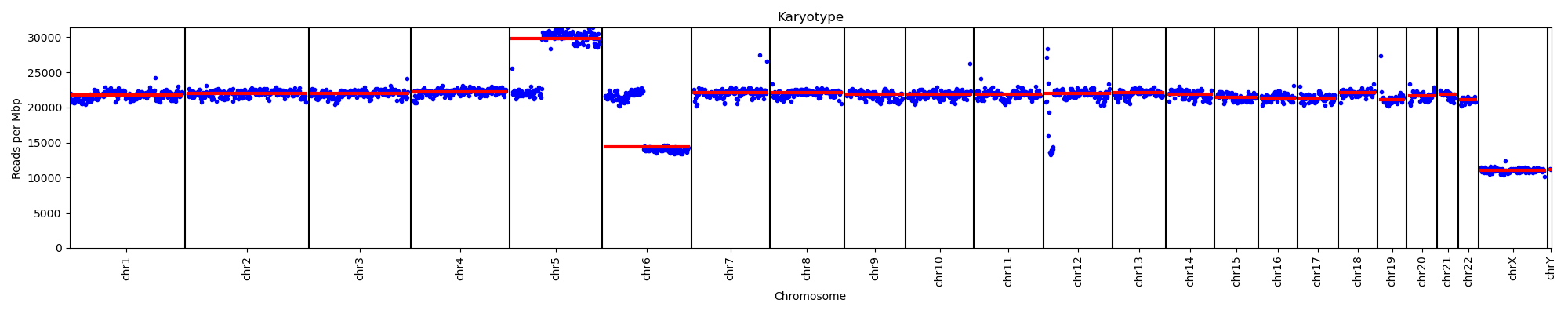

Nano_Leuk_0244
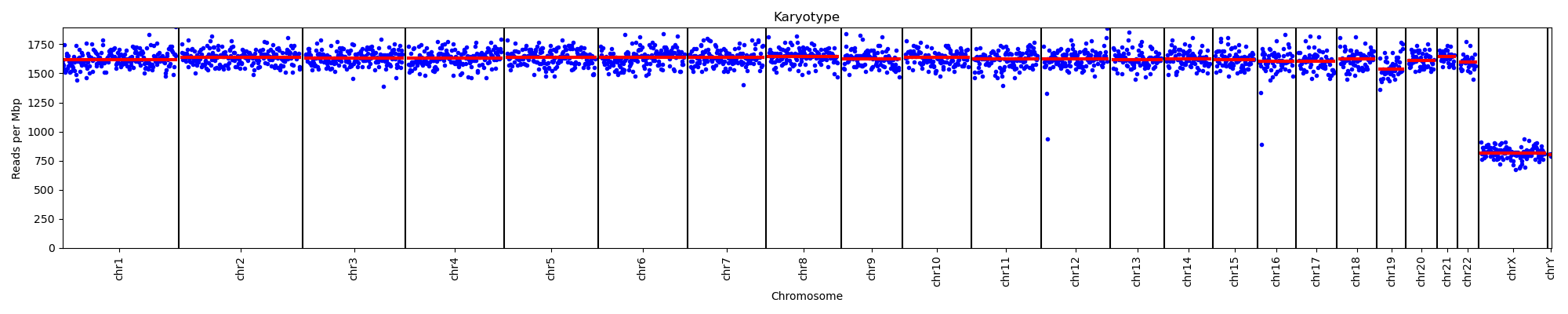

Nano_Leuk_0250
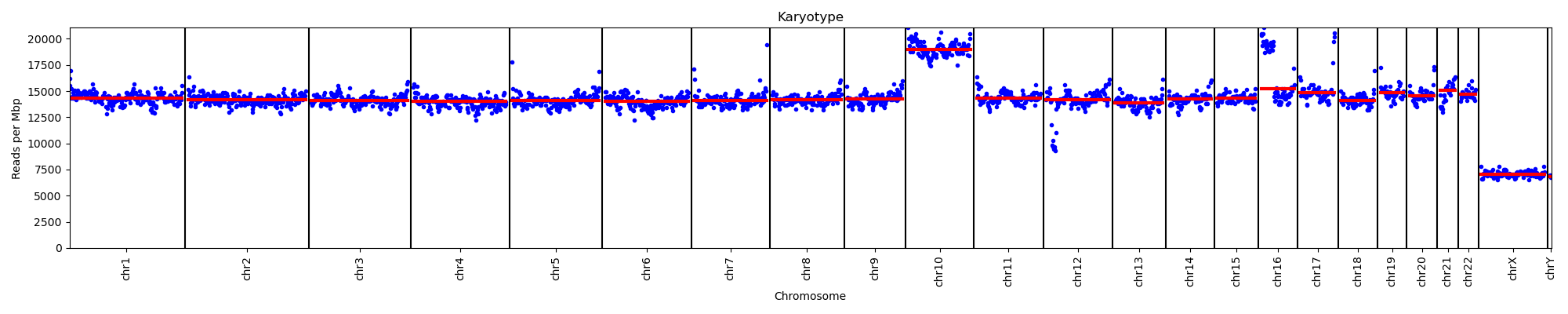

Nano_Leuk_0254

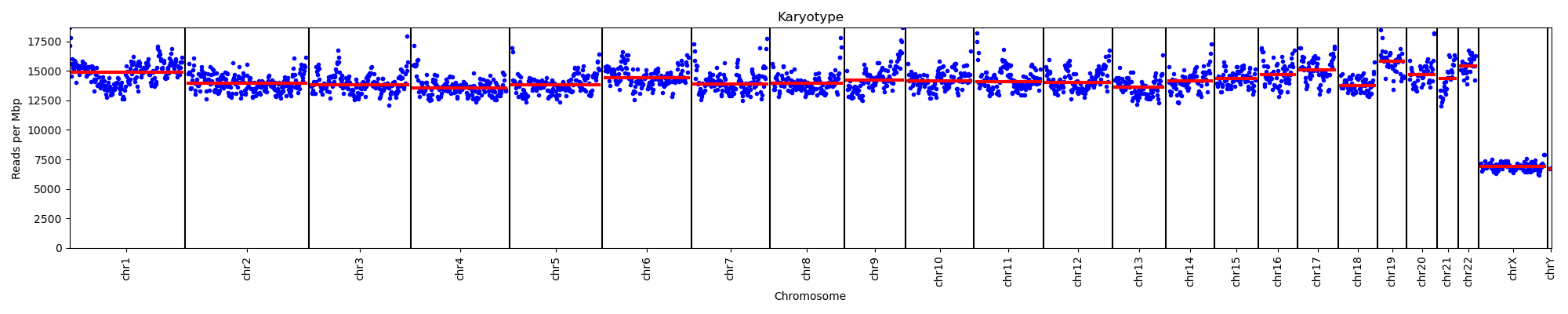
